# *Mycobacterium tuberculosis* MutT4 is an RNA pyrophosphohydrolase that forms biomolecular condensates and sensitizes mRNAs to degradation

**DOI:** 10.1101/2025.05.13.653832

**Authors:** J. Hilario Cafiero, Junpei Xiao, Irene Lepori, Abigail R. Rapiejko, Manchi Reddy, Louis A. Roberts, M. Sloan Siegrist, James C. Sacchettini, Scarlet S. Shell

## Abstract

Bacterial adaptation to stress involves changes in transcription and mRNA degradation rates. In *Escherichia coli*, the Nudix hydrolase RppH initiates mRNA degradation by removing pyrophosphate from mRNA 5’-ends, converting 5’-triphosphates to 5’-monophosphates. We aimed to identify the RppH homolog in the globally important pathogen *Mycobacterium tuberculosis* (Mtb). We deleted each non-essential Nudix gene from Mtb to determine their impacts on mRNA phosphorylation states. Deletion of *mutT4* (Rv3908) increased the relative abundance of 5’-triphosphates on myriad mRNAs across the transcriptome. Purified MutT4 converted mRNA 5’-triphosphates into monophosphates, and stimulated degradation by RNase E and RNase J. MutT4 has intrinsically disordered regions (IDRs), a common domain for biomolecular condensate formation. Microscopy showed that MutT4 forms condensates that dissociate upon addition of rifampicin, and that the N-terminal IDR is sufficient for condensate formation. These MutT4 condensates localize with RNase E and RNase J. Deletion of *mutT4* in Mtb leads to a higher outer membrane permeability and resistance to oxidative stress. We conclude that MutT4 is the RppH homolog of Mtb, assembling in condensates that may act as degradation hubs. Our data indicate that MutT4 is unlikely to participate in DNA repair or nucleotide pool cleansing, and as such would more accurately be named RppH.

## INTRODUCTION

*Mycobacterium tuberculosis* (Mtb), the causative agent of tuberculosis (TB), has reclaimed its position as the leading cause of death from a single infectious agent, surpassing SARS-CoV-2 (1). In 2023 alone there were 8.2 million reported TB cases and 1.25 million estimated deaths (1). Standard treatment requires a 4-to 6-month course of multiple antibiotics. Mtb’s success as a pathogen and recalcitrance to treatment are due in part to its abilities to overcome the plethora of stressors it encounters within the host, such as reactive oxygen and nitrogen species, transition metal stress (Cu^+^, Zn^2+^), low pH, and hypoxia, among others (reviewed in (2)).

Bacterial adaptation to stress involves changes in RNA expression levels. The steady-state abundance of any given RNA results from the balance between its rate of transcription by RNA polymerase and its rate of degradation by RNases and auxiliary proteins such as helicases. mRNA degradation is a regulated process, but the mechanisms underlying its regulation in mycobacteria are not well understood. Severe energy stressors such as hypoxia and carbon starvation lead to global slowing of mRNA degradation rates in mycobacteria through largely unknown mechanisms (3–5). Meanwhile, translation and stability of some specific transcripts can be regulated by small RNAs (6–13). Developing a comprehensive understanding of the mechanisms governing regulation of mRNA degradation is hampered in part by gaps in knowledge about the machinery that carries out mRNA degradation. Mycobacteria encode a number of RNases, including the essential RNase E which has a rate-limiting step in degradation of most mRNAs in *Mycolicibacterium smegmatis* (a non-pathogenic model mycobacterium) (14) and RNase J which has a specialized role in mRNA degradation and affects drug sensitivity in Mtb (15). The physical and functional interactions among these and other enzymes are largely undefined in mycobacteria. In *Escherichia coli* and some other bacteria, complexes of RNases called degradosomes have been defined and appear to increase the efficiency of mRNA degradation (16,17). In Mtb, some RNases and putative RNA binding proteins were shown to co-precipitate, suggesting multiple possible degradosome structures, although the direct protein-protein interactions mediating these remain mostly undefined (18). Interestingly, degradosome composition in *E. coli* and *Caulobacter crescentus* seems to vary in response to microenvironmental conditions (19,20), and RNA degradation proteins in *Bacillus subtilis* were proposed to interact in a transient degradosome-like network rather than as a stable complex with defined stoichiometry (21). Recent evidence shows that in *C. crescentus*, the enzymes that form RNA degradosomes also assemble into large, dynamic protein condensates within the bacterial cytoplasm, in a process that involves liquid-liquid phase separation (22). mRNA degradation therefore appears to be regulated at multiple levels including specific protein-protein interactions, weak multivalent interactions, and subcellular localization.

Some clinical strains of Mtb carry drug resistance-associated mutations in RNA processing and degradation enzymes (23). For instance, loss of RNase J in Mtb increases drug tolerance by altering mRNA metabolism and impacting gene expression (15). Although mRNA degradation pathways in Mtb are linked to drug resistance and have been proposed as targets for novel therapeutics (24), there is a key step in RNA degradation that has not yet been described in mycobacteria. Transcription of most bacterial mRNAs initiates with nucleoside triphosphates and primary transcripts therefore have triphosphates on their 5’ ends. In *E. coli* and *B. subtilis*, these 5’ triphosphates appear to have a role analogous to 5’ caps on eukaryotic mRNAs; conversion of 5’ triphosphates to 5’ monophosphates by RNA pyrophosphohydrolase (RppH) sensitizes mRNAs to degradation (25,26). 5’ monophosphates stimulate the endonuclease activity of *E. coli* RNase E by an allosteric interaction (27,28), while for RNase J, exonuclease activity is stimulated by these substrates (26,29–32). Pyrophosphate removal by RppH has been demonstrated as a rate-limiting step in RNA degradation for a subset of transcripts in *E. coli* (25,33), *B. subtilis* (26,34) and *Helicobacter pylori* (35). No RppH ortholog has been identified or annotated in mycobacteria. However, 5’-end-directed RNAseq showed the presence of 5′ monophosphate-bearing transcripts in both Mtb and *M. smegmatis* (14), indicating that there is a native enzyme in mycobacteria that converts RNA 5’ triphosphates into monophosphates. Here we show that MutT4 (Rv3908), previously thought to participate in nucleotide pool cleansing (36), is actually the RppH ortholog of Mtb. We show for the first time that conversion of 5’ triphosphates to monophosphates stimulates cleavage by Mtb RNase E and RNase J. Unlike the RppHs of *E. coli* and *B. subtilis*, mycobacterial MutT4 has intrinsically disordered regions (IDRs) that trigger formation of biomolecular condensates in the bacterial cytoplasm. Finally, deletion of *mutT4* affects various clinically important phenotypes in Mtb by simultaneously increasing outer membrane permeability and resistance to oxidative stress.

## MATERIALS AND METHODS

### Bacterial strains and growth conditions

*M. tuberculosis* mc^2^6030 H37Rv Δ*RD1* Δ*panCD* (37) and derivative strains were cultured in Middlebrook 7H9 supplemented with OADC (0.05 g/L oleic acid, 5 g/L bovine serum albumin fraction V, 2 g/L glucose, 0.85 g/L NaCl, and 4 mg/L catalase), 0.2% glycerol, 0.05% Tween 80, and 24 μg/mL pantothenate at 37 °C and 200 rpm shaking. Middlebrook 7H10 supplemented with 0.5% glycerol, OADC, and 24 μg/mL pantothenate was used for growth on solid media. *Mycolicibacterium smegmatis* mc^2^155 (Msmeg) was grown in the same conditions but oleic acid and pantothenate were excluded from the media (7H9 DC). Apramycin (5 μg/mL), hygromycin (50 μg/mL for Mtb, 150 μg/mL for Msmeg), or zeocin (25 μg/mL) were added to the media when required. The recombineering system described in (38) was used to generate strains deleted for each of ten non-essential genes predicted to encode Nudix proteins. Mutants were generated by replacing most of the coding sequence of each gene with a zeocin resistance gene under the EM7 promoter. The first 10-51 codons of each gene were left intact in order to prevent disruption of overlapping or divergently transcribed genes. These codons were followed by a stop codon, the EM7 promoter, 5’ UTR, zeocin coding sequence, and 4-105 codons at the end of the Nudix gene as needed to prevent disruption of expression of downstream genes. Sequencing of PCR amplification products with primers flanking the deleted region of each strain was performed to confirm the expected sequence. A Giles-site integrating plasmid containing *mutT4* and its native promoter and 5’ UTR (560 bp upstream of the start codon and 300 bp upstream of the TSS) was used for complementation. All DNA constructs were made with NEBuilder HiFi DNA Assembly (New England Biolabs, E2621). Primers, plasmids and strains used in this study are listed in **Table S1**.

### RNA purification

Mtb cultures in exponential phase (OD_600nm_ = 0.8 – 1.6) were centrifuged at 3,900 rpm for 7 min at 4 °C. The bacterial pellets were resuspended in 1 mL of TRI Reagent (Molecular Research Center, TR118) and transferred to bead-beating tubes (OPS Diagnostics 100 µm zirconium beads, PFMB 100-100-12). Cells were lysed with 3 cycles at 7 m/s, 30 seconds per cycle in a FastPrep-24 bead-beater (MP Biomedicals) and then 300 µL of chloroform was added. Samples were centrifuged 15,000 rpm for 15 min at 4 °C and RNA from the aqueous phase was purified using Direct-Zol RNA miniprep kit (Zymo Research, R2052) following the manufacturer’s instructions. DNase treatment was performed on-column and the purified RNA was stored at -80 °C.

### Splinted ligation

Splinted ligation was performed to quantify the relative amounts of RNA 5’ tri and diphosphates (henceforth referred to collectively as triphosphates, because they cannot be distinguished from each other by this method) or 5’ monophosphates following the method described in (39) with modifications (**Figure 1 A**). 5 µg of DNase-treated total RNA extracted from Mtb in exponential phase was incubated with 10 U of *E. coli* RNA 5’ Pyrophosphohydrolase (RppH) (New England Biolabs, M0356), or with no enzyme as a control, in NEBuffer 2 for 1 h at 37 °C. RNA was purified with RNA Clean & Concentrator-5 (Zymo Research, R1014) following the manufacturer’s instructions. 1 µg of RppH-treated RNA (or no enzyme control) was mixed with a chimeric DNA/RNA adapter oligonucleotide (SSS1016) and a DNA oligo splint (SSS3118) targeting the previously mapped +1 nucleotide of Rv3248c mRNA (14,40) and incubated for 5 minutes each at 70 °C, 60 °C, 42 °C, and 25 °C. Next, 1,000 U of T4 DNA Ligase (New England Biolabs, M0202), 40 U of RNase Inhibitor, Murine (New England Biolabs, M0314) and T4 DNA ligase buffer to a final 1X concentration were added and the sample was incubated at 15 °C overnight. Ligation occurs at this step only in Rv3248c mRNA with a monophosphate at its 5’-end. The splint increases the efficiency of ligation to the exact 5’ end of the Rv3248c mRNA. RNA was purified with RNA Clean & Concentrator-5 and used for cDNA synthesis as follows. RNA was incubated 10 min at 70 °C with 1.6 µM of a gene-specific reverse primer for Rv3248c (SSS2650) in Tris HCl 12.8 mM pH 7.5. Then, the sample was mixed with 100 U of ProtoScript II Reverse Transcriptase (New England Biolabs, M0368), 10 U of RNase Inhibitor Murine, 0.5 mM dNTPs, 5 mM DTT and 1X ProtoScript II Reverse Transcriptase Reaction Buffer. The sample was incubated for 10 min at 25 °C and 2 h at 42 °C for cDNA synthesis. Quantitative PCR (qPCR) with iTaq Universal SYBR Green Supermix (Bio-Rad, 1725124) was performed to quantify two amplicons derived from the cDNA: one internal amplicon of the Rv3248c gene to use for normalization, and one amplicon derived from the junction of the adapter and Rv3248c ligation product. qPCR was performed in a QuantStudio 6 Pro (Applied Biosystems, ThermoFisher Scientific) thermocycler with 0.25 μM of each primer (**Table S1 C**) in 10 μL reaction mixtures, with a cycle consisting of: 2 min at 50 °C, 10 min at 95 °C followed by 40 cycles of 15 s at 95 °C and 1 min at 60 °C. All primers used for qPCR had an amplification efficiency between 90 and 110% as determined by linear regression of serial dilution of cDNA samples.

**Figure 1.**
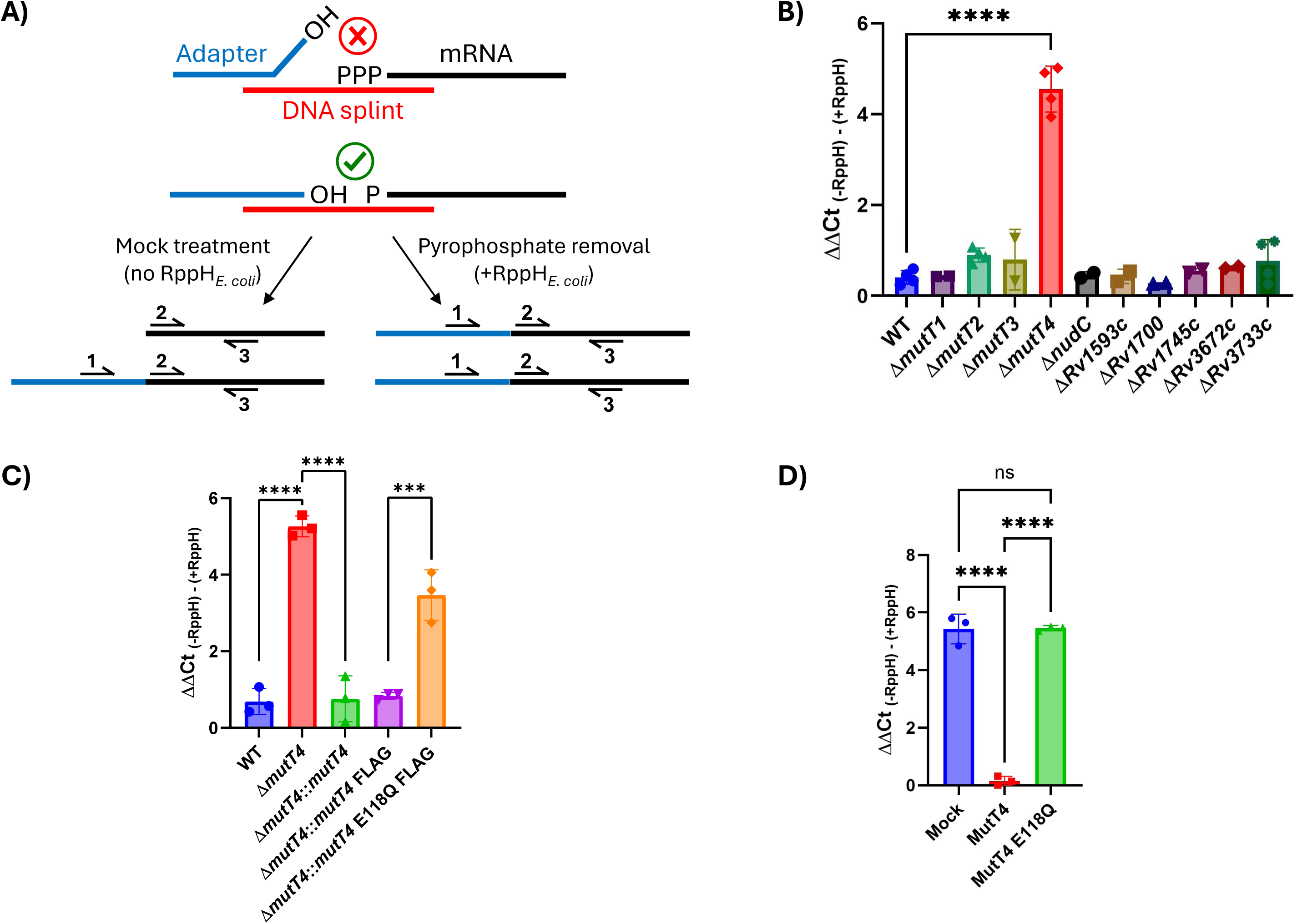
MutT4 is an RNA pyrophosphohydrolase that acts on Mtb mRNAs. **(A)** Schematic of splinted ligation. The adapter is ligated only if the target mRNA has a 5’ monophosphate. Ligation is performed with and without treatment by *E. coli* RppH to convert 5’ triphosphates to monophosphates. Quantitative PCR reveals the abundance of ligated mRNA (primers 1 and 3) relative to total mRNA (primers 2 and 3). **(B-C)** Screening of Mtb Nudix gene deletion mutants and complemented strains by splinted ligation of a transcript in the endogenous mRNA pool. ΔΔCt indicates the impact of in vitro treatment with *E. coli* RppH on the ligatability of the Rv3248c transcript, after normalizing to total transcript abundance. Higher ΔΔCt values indicate a higher proportion of the endogenous Rv3248c mRNA was triphosphorylated. Mean and standard deviation of three biological replicates are shown for panel C. **(D)** MutT4 has RNA pyrophosphohydrolase activity *in vitro*. Splinted ligation of *in vitro* transcribed Rv3248c mRNA (5’ monophosphorylated) treated with purified MutT4, MutT4 E118Q, or no enzyme (mock). Each value represents results from the assay performed with an independent protein preparation. A one-way ANOVA was performed with Dunnett’s test for panel B and Tukey’s test for panel C and D. *** p ≤ 0.001, **** p ≤ 0.0001.

### RNAseq

RNA was extracted from Mtb exponential growth phase cultures as described above. rRNA was depleted as stated in (41). 5’-end-directed libraries were constructed as stated in (42) with the following modifications. rRNA was depleted as stated above. For normalization, 2.5 pg of *in vitro* transcripts (IVTs) (see below) were added at the adapter ligation step. Fragmentation was performed for 15 min at 94 °C. PCR primers for adding full-length Illumina adapter sequences are listed in **Table S1 C**, and the primers used for each library are listed in **Table S1D**. Selection of library fragments was done with a double-sided purification using HighPrep PCR Clean Up System (MagBio, AC-60005). Libraries were pooled and sequenced at SeqCenter (Pittsburgh, PA).

IVTs used as spike ins for normalization were designed to include synthetic transcripts absent in Mtb (**Table S2**). IVTs were synthesized by *in vitro* transcription with HiScribe T7 Quick High Yield RNA Synthesis Kit (New England Biolabs, E2050) using as templates purified PCR products containing a T7 promoter and the coding sequence of each transcript. IVT RNA products were separated in a 1% agarose TBE gel and bands corresponding to the expected transcript size were excised and purified using the Zymoclean Gel RNA Recovery Kit (Zymo Research, R1011). IVTs were mixed in equimolar amounts and treated with 5 U of RppH (New England Biolabs) for 1 h at 37 °C in NEBuffer 2 and purified with RNA Clean & Concentrator-5 (Zymo Research).

### Bioinformatics tools and analyses

Reads from Mtb 5’-end-directed libraires were aligned to the NC_000962 reference genome using Burrows-Wheeler Aligner (43). Due to the library construction method, Read 1 corresponds to the sequence of an endogenous RNA, with its first nucleotide representing the 5’-most nucleotide of that RNA molecule, and Read 2 corresponds to the reverse complement of part of an RNA. Samtools was used to remove Read 2 and separate Read 1s that mapped to plus and minus strands of the genome (44). Bedtool2 was used to obtain the read depth for each coordinate at the 5’ end of a Read 1, representing the 5’ end of an RNA used to construct the library (45). The total number of reads mapped to spike-in controls in each library was used to normalize the coordinate-based coverage of 5’ ends. To identify biologically relevant 5’ ends, those with a normalized coverage below 5 were excluded. The log₂ ratio of the filtered 5’-end coverage in +RppH/-RppH conditions was then calculated for each strain. Only the 5’ ends that were previously annotated as transcription start sites (TSSs) and cleavage sites were used for downstream analysis (14). Normalized coverage at each of these coordinates in each library is reported in Tables S4 and S5. The distribution of the log₂ ratio of 5’-ends in +RppH/-RppH is expected to be bimodal, with one population centered around zero and another with higher log₂ ratios, representing TSSs (42). The 20-nucleotide sequence, starting from the 5’-end (the first nucleotide), was extracted and used to compute the minimum free energy (MFE) and unpaired probability using RNAfold (46). Gene cluster comparison figures were created with Clinker (47). Intrinsically disordered protein regions were analyzed with IUPred3 (48).

### Expression and purification of MutT4

*Escherichia coli* BL21(DE3) pLysS transformed with pET42 *mutT4* or pET42 *mutT4* E118Q (catalytically inactive mutant) were grown at 37 °C with 200 rpm shaking in LB with 50 µg/mL kanamycin and 20 µg/mL chloramphenicol to OD_600nm_ = 0.6. IPTG was added at 0.1 mM and bacteria were incubated at 30 °C 200 rpm for 3 hours to induce protein expression. The culture was centrifuged, and the bacterial pellet from 1 L of culture was resuspended in 10 mL equilibration buffer (Tris HCl 20 mM pH 8, NaCl 300 mM, glycerol 5% (v/v), IGEPAL 0.01% (v/v), imidazole 10 mM), supplemented with 20 U of Turbo DNase (ThermoFisher Scientific, AM2238) and 1.25 mg/ml of lysozyme (ThermoFisher Scientific, J60701.06). The sample was transferred to 100 µm zirconium beads tubes (∼1.5 mL/ tube; OPS Diagnostics, PFMB 100-100-12) and cells were lysed with a FastPrep-24 bead-beater (MP Biomedicals) using 4 cycles of 30 sec at 6.5 m/s. Lysates were centrifuged 20 min 13,000 rpm at 4 °C and the supernatant was incubated with HisPur Ni-NTA resin (ThermoFisher Scientific, 88221) and Halt Protease Inhibitor Cocktail 1X (ThermoFisher Scientific, 78430) for 1 hour at room temperature with end-to-end rotation. The resin was washed six times with wash buffer (Tris HCl 20 mM pH 8, NaCl 1 M, glycerol 5% (v/v), IGEPAL 0.01% (v/v), imidazole 50 mM) and then elution buffer (Tris HCl 20 mM pH 8, NaCl 300 mM, glycerol 5% (v/v), IGEPAL 0.01% (v/v), imidazole 500 mM) was added to elute the purified protein. The eluted samples were dialyzed in Slide-A-Lyzer 20,000 MWCO cassettes (ThermoFisher Scientific, 66003) against storage buffer (Tris HCl 20 mM pH 8, NaCl 300 mM, glycerol 5% (v/v), IGEPAL 0.01% (v/v)). Purified protein samples were flash frozen in liquid N_2_ and stored at -80 °C.

### MutT4 enzymatic characterization

Rv3248c mRNA was synthesized by *in vitro* transcription (IVT) with HiScribe T7 Quick High Yield RNA Synthesis Kit (New England Biolabs, E2050) using as a template a purified PCR product (amplified with primers SSS3829 and SSS3830) containing a T7 promoter and the first 359 bp of Rv3248c mRNA starting from the +1 transcription start site. The IVT RNA was purified using RNA Clean & Concentrator-25 (Zymo Research, R1018). A single RNA band was confirmed by running an aliquot of the purified RNA on a 1% agarose TBE 1X gel. 0.5 µg of purified MutT4 or MutT4 E118Q were mixed with 250 ng of Rv3248c mRNA in a buffer containing Tris HCl 40 mM pH 8, MgCl_2_ 10 mM, DTT 2 mM, glycerol 2% (v/v), NaCl 600 mM and incubated for 1 hour at 37 °C. As a control, samples were incubated without the addition of protein. RNA was purified with RNA Clean & Concentrator-5 and the triphosphate to monophosphate ratio at the RNA 5’-end was measured with splinted ligation as stated previously.

### Expression and purification of RNase E and RNase J

Mtb RNase E (Rv2444c) and RNase J (Rv2752c) were cloned into pET vectors and overexpressed in *E. coli* with C-terminal 6xHis tags (complete tag sequence: LEHHHHHH). The proteins were purified over Ni-NTA resin followed by size exclusion chromatography. Briefly, *E. coli* pellets were lysed in 20 mM Tris-HCl (pH 7.5), 500 mM NaCl, and 5% glycerol, loaded on Ni-NTA resin. The resin was washed with buffer containing 100 mM imidazole and 1 M NaCl, then eluted in a gradient from 75 to 500 mM imidazole. Proteins were then further purified with Superdex size-exclusion column in buffer containing 50 mM Tris-HCl, pH 7.5, 100 mM NaCl, 5% glycerol, and 2 mM DTT.

### RNA cleavage assay

Rv3248c mRNA was synthesized and treated with MutT4 as stated above. RNA was purified after treatment with RNA Clean & Concentrator-5. For each reaction 40 ng of RNA were incubated for 3 min at 65 °C. Next, the RNA was diluted in a buffer containing Tris HCl 20 mM pH 8.0, DTT 1 mM, IGEPAL 0.01% (v/v), glycerol 0.5% (v/v), MgCl_2_ 10 mM, ZnCl_2_ 10 µM, NaCl 150 mM, and 80 ng of purified Mtb RNase E, Mtb RNase J, or no enzyme added as a control in 10 µl of final volume. Samples were incubated for 30 min at 37 °C. The reaction was stopped by addition of 1X RNA gel loading dye (Thermo Scientific, R0641) and incubation for 10 min at 70 °C. Samples were analyzed in a 1X TBE Urea PAGE gel stained with SYBR Gold (Thermo Scientific, S11494). Quantification of band intensity was performed with ImageJ with Fiji plugin (49). Cleavage was expressed as percentage decrease in substrate band intensity after incubation with RNase E or RNase J compared to the corresponding untreated control.

### Western blotting

Mtb samples grown to exponential phase were centrifuged for 7 min 3,900 rpm at 4 °C and washed with TBS (Tris HCl 20 mM pH 7.6, NaCl 150 mM). Bacterial pellets were resuspended in lysis buffer (HEPES 50 mM pH 7.5, NaCl 150 mM, EDTA 1 mM, Triton X-100 0.5% (v/v), glycerol 10% (v/v), SDS 1% (w/v)), and transferred to tubes containing 100 µm zirconium beads (OPS Diagnostics, PFMB 100-100-12). Cells were lysed with a FastPrep-24 bead-beater (MP Biomedicals) using 4 cycles of 30 sec at 6.5 m/s. Lysates were centrifuged for 10 min 13,000 rpm at 4 °C and the supernatants were transferred to new tubes. Protein concentration in the lysates was quantified using Pierce BCA Protein Assay Kit (ThermoFisher Scientific, 23227). Equal amounts of protein from each sample were separated on SDS PAGE gels and transferred to PVDF membranes. Membranes were stained with LiCor Revert 700 Total Protein Stain (LiCor, 926-11021) following the manufacturer’s instructions and imaged to verify equal loading. The membrane was blocked with TBS skim milk 3% for 45 min at room temperature, washed with TBS and incubated with monoclonal anti-FLAG M2 mouse antibody (1 µg/mL) (Sigma-Aldrich, F1804) in TBS skim milk 3% overnight at 4 °C. Membranes were washed with TBS, and then incubated with goat anti-mouse HRP antibody for 45 min at room temperature. Membranes were washed with TBS Tween-20 0.05% (v/v) and then developed with Azure Radiance ECL (AC2204) in an Azure 600 imager.

### Microscopy

Microscopy images were acquired with a Nikon CSU W1 spinning disc confocal microscope equipped with a CFI 60 plan apochromat λ D 100× Oil immersion objective. *M. smegmatis* cultures in exponential phase were washed once in 7H9 without DC and then spotted onto 1.2% agarose pads and imaged in a chamber at 37 °C. Dendra2 was imaged using a 488 nm excitation laser and a ET525/36m emission filter. For mScarlet a 561 nm excitation laser and a ET605/52m emission filter were used. For rifampicin treatment, exponential phase cultures were incubated with rifampicin 100 µg/mL at 37 °C and 200 rpm, and samples were taken after 30 min for imaging. A sample with DMSO (rifampicin vehicle) was included as a no drug control. Images were acquired, denoised, and deconvoluted using Nikon Elements software (5.42.06). Image analysis was performed using ImageJ with Fiji plugin. The number of condensates per bacteria was quantified as the coefficient of variation, as an unbiased measurement independent of cell length.

### Antibiotic susceptibility testing

Susceptibility to antibiotics was determined using a resazurin microtiter assay. Mtb was grown to exponential phase and inoculated in 96 well plates with different concentrations of antibiotics in 200 µL of final volume and a bacterial OD_600nm_ = 0.01. Plates were incubated for 10 days at 37 °C and 125 rpm shaking. Resazurin 0.002% (w/v) was added to each well, incubated for 24 h and absorbance of reduced resazurin was measured at 570 nm. The bacterial growth of each well containing antibiotic was normalized to growth in a control well without antibiotic.

### Hypoxic growth conditions

A variation of the Wayne model (50) was used to determine the bacterial viability in hypoxia as described in (51). Vials were opened after 3 or 6 weeks, and cultures were diluted and plated on 7H10 to enumerate CFUs.

### Resistance to hydrogen peroxide

A disk diffusion assay was used to measure the resistance to H_2_O_2_. Mtb was grown to exponential phase in 7H9 without the addition of catalase (7H9 OAD), diluted to OD_600nm_ = 0.01 in 7H9 OAD containing agar 0.6 % (w/v) (12 mL final volume), and plated on top of 25 mL 7H10 OAD in a 100 mm round Petri dish. Paper disks with 10 µl of 3% H_2_O_2_ were added on top of the solidified agar. Plates were incubated for 10 days at 37 °C and the inhibition zone was measured. For acute H_2_O_2_ treatment, Mtb was grown to exponential phase in 7H9 OAD, diluted to OD = 0.1 and H_2_O_2_ was added to a final concentration of 10 mM. Cultures were incubated for 24 h at 37 °C with 120 rpm shaking, and serial dilutions were plated on 7H10 to enumerate CFUs. Survival was expressed as the percentage of CFUs recovered at 24 h compared to before the addition of H_2_O_2_.

### Permeability assay (PAC-MAN) of azide-rifamycin on live *M. tuberculosis*

To evaluate the permeability of mycomembrane to rifamycin we perform the previously described <u>P</u>eptidoglycan <u>A</u>ccessibility <u>C</u>lick-<u>M</u>ediated <u>A</u>ssessme<u>N</u>t (PAC-MAN) assay (52–55). Mtb wild type, Δ*mutT4*, and the complemented strain Δ*mutT4::mutT4* were diluted to OD_600_ ∼0.04 and grown in 7H9 +/- 25 μM dibenzocyclooctyne (DBCO)-tetrapeptide (TetD (52) WuXi AppTec) for 3 days until OD_600_ reached ∼0.3-0.4. Bacteria were washed twice with polyphosphate buffer supplemented with Tween-80 0.05% (PBST) then incubated with 50 μM of azide-rifamycin (56) in PBST in 96-well plates for 2 hours at 37 °C with shaking. Azide-rifamycin was removed by spinning, the supernatant was removed, and bacteria were then incubated in PBST containing 17 μM of the fluorogenic label CalFluor647-azide (Vector laboratories, APC flow cytometry channel) for 1 hour at 37 °C with shaking. The fluorophore was removed by spinning and supernatant removal and bacteria were fixed with fresh prepared 4% paraformaldehyde (PFA, Ted Pella) in PBS for 2 hours at room temperature. Finally, bacteria were resuspended in physiological solution (0.9% NaCl in water) and analyzed by BD DUAL LSRFortessa flow cytometer.

### Fluorescent-vancomycin permeability assay

Mtb wild type, Δ*mutT4*, and the complemented strain Δ*mutT4::mutT4* were grown to OD_600_ ∼0.1, concentrated to a working OD_600_ of 0.4 in PBS, and incubated with BODIPY-vancomycin (FL-VAN, Thermo Fisher, FITC flow cytometry channel) 1:1 with native vancomycin, both 5 μg/mL, for 4 h in PBS at 37°C with shaking (57). Bacteria were then washed 3 times with PBST, fixed with fresh prepared 4% PFA for 2 hours at room temperature, resuspended in physiological solution and analyzed by BD DUAL LSRFortessa flow cytometer. Fluorescence measurements were normalized to the wild type strain.

## RESULTS

### MutT4 is the Mtb RNA pyrophosphohydrolase

All the RNA pyrophosphohydrolases described so far in bacteria belong to the Nudix hydrolase family (25,26,35,58). We hypothesized that the RppH ortholog of Mtb also belongs to this protein family. The Mtb H37Rv genome has 11 genes with Nudix domains, only one of which is essential (**Supplementary Table 3**). To identify the Nudix protein that converts RNA 5’ triphosphates to monophosphates in Mtb, we generated strains with single deletions of each non-essential Nudix gene. Next, we screened the mutant collection by splinted ligation to quantify the 5’-end phosphorylation status of the mRNA target Rv3248c. We chose this target because in the WT background it is mainly monophosphorylated (14), suggesting that it is targeted by the native RNA pyrophosphohydrolase. The splinted ligation method compares the relative abundance of adapter-ligated RNA with and without treatment of the RNA pool with *E. coli* RppH (**Figure 1 A**). Because ligation requires a 5’ monophosphate, RNAs not treated with RppH are only ligated if they have 5’ monophosphates when extracted from Mtb. Deletion of *mutT4* (Rv3908) significantly increased the 5’-end triphosphates of the target mRNA as compared to the WT strain (**Figure 1 B**), suggesting that in the WT strain *mutT4* is involved in conversion of 5’ triphosphates to monophosphates on this transcript. Integration of a single copy of *mutT4* under its native promoter in the Δ*mutT4* mutant strain reverted the 5’-end triphosphates of target Rv3248c to WT levels (**Figure 1 C**). Analysis of gene expression by qPCR showed that the WT and complemented strains expressed *mutT4* at similar levels (**Supplementary Figure 1**). *mutT4* is conserved among the mycobacteria, and it is coded downstream of *pcnA*, recently described as the tRNA 3’-end CCA-adding enzyme of Mtb (59), indicating that this genomic region codes for RNA modifying enzymes (**Supplementary Figure 2**).

To confirm the enzymatic activity of MutT4, we complemented the Δ*mutT4* mutant strain with a *mutT4* variant harboring a point mutation in the predicted catalytic site in the Nudix domain to abrogate its enzymatic function (Glu 188 to Gln, E188Q). Point mutations in this key residue have been reported to inactivate the catalytic activity of RppH in *E. coli*, *B. subtilis*, and *H. pylori* (25,26,35). Western blotting revealed that FLAG-tagged WT and E188Q MutT4 had equivalent protein abundance in complemented strains (**Supplementary Figure 3**). However, complementation of the Δ*mutT4* mutant strain with the E118Q allele failed to revert the 5’-end phosphorylation status of Rv3248c to WT levels (**Figure 1 C**), suggesting that the Nudix motif is required for the pyrophosphohydrolase activity. To further confirm that MutT4 acts directly upon mRNAs, we expressed and purified Mtb MutT4 protein (either WT or E118Q) from *E. coli* and tested its RNA pyrophosphohydrolase activity *in vitro* using as a substrate Rv3248c mRNA synthetized by *in vitro* transcription with 5’ triphosphates. Incubation with MutT4^WT^ significantly converted the 5’-end triphosphates to monophosphates as compared to a control where the mRNA was incubated without enzyme (**Figure 1 D**). No reduction in 5’-end triphosphates was observed when incubating the RNA with the catalytically-dead MutT4^E118Q^ (**Figure 1 D**), indicating that this activity was not from a contamination present in the protein preparation. These results show that MutT4 is a *bona fide* RNA pyrophosphohydrolase.

### MutT4 acts on Mtb mRNAs transcriptome-wide

To determine which mRNAs are MutT4 substrates transcriptome-wide in Mtb, we next performed 5’-end-directed RNAseq (**Figure 2 A**). This method compares the abundance of adapter-ligated RNA 5’ ends with and without treatment of the RNA pool with *E. coli* RppH *in vitro* prior to ligation and library construction. The difference in abundance of a given RNA 5’ end with and without RppH treatment therefore reflects the extent to which it was triphosphorylated prior to extraction from Mtb. We examined the relative phosphorylation states of RNA 5’ ends corresponding to transcription start sites (TSSs) previously mapped in Mtb (14,40) (**Table S4**). Deletion of *mutT4* dramatically increased the relative abundance of 5’ triphosphates on myriad transcripts (**Figure 2 B**), and these were reverted to WT levels by complementation. This result shows that MutT4 has a transcriptome-wide role in converting RNA 5’ triphosphates to monophosphates. We also examined 5’-end reads corresponding to previously mapped RNase cleavage sites as a normalization control, since cleavage by the RNases most active in Mtb produces 5’ monophosphates (14). As these reads derived from exclusively monophosphorylated 5’-ends, treatment with RppH should not increase their coverage ratio, which is what we observed (**Figure 2 B, Table S5**).

**Figure 2.**
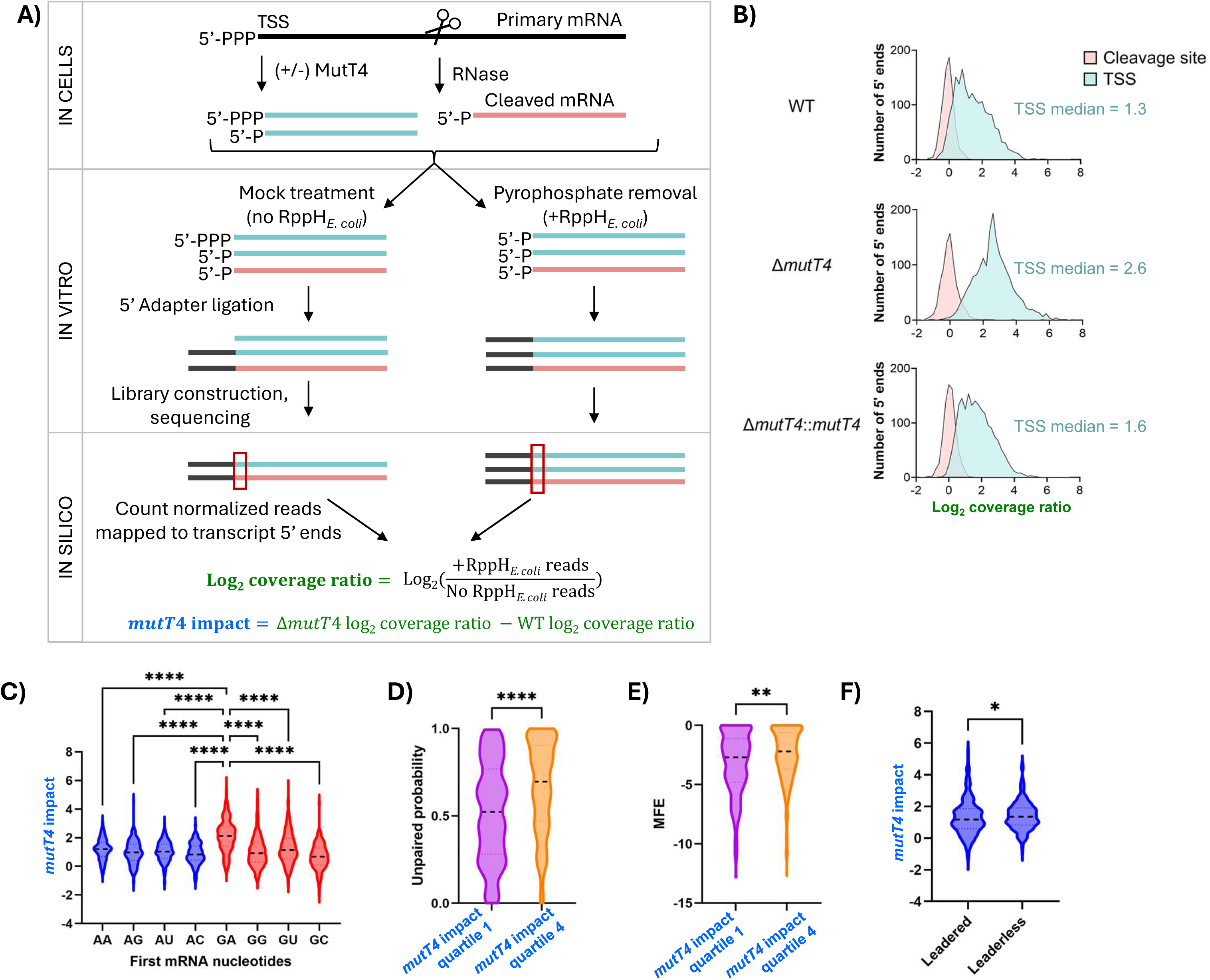
MutT4 has a transcriptome-wide role as an RNA pyrophosphohydrolase, with a preference for transcripts that start with guanosine and have minimal secondary structure near their 5’ ends. **(A)** Schematic of 5’-end directed RNASeq. Total RNA was used to construct 5’-end-directed libraries with or without pre-treatment with *E. coli* RppH to convert 5’ triphosphates to monophosphates. Only transcripts with 5’ monophosphates (whether endogenous or produced by *E. coli* RppH) can be ligated to adapters and therefore be captured in the libraries. Comparison of the number of reads obtained in libraries made from RppH-treated vs untreated RNA indicates the extent to which the RNA was endogenously monophosphorylated. The “log_2_ coverage ratio” (green) and “*mutT4* impact” (blue) are metrics used in subsequent panels. **(B)** Log_2_ coverage ratio of +/-RppH libraries for reads derived from RNA 5’ ends previously classified as transcription start sites (TSS, teal curve, n=1687) or RNase cleavage sites (pink, n=590) (Zhou and Sun et al 2023). The RNase cleavage sites produce RNAs with monophosphorylated 5’ ends and are therefore expected to have log_2_ coverage ratios of ∼0, while TSSs produce RNAs with triphosphorylated 5’ ends and are therefore expected to have log_2_ coverage ratios >0. The median TSS log_2_ coverage ratio for each strain is shown in teal. **(C-F)** For each TSS quantified in panel B, the impact of *mutT4* on 5’ end phosphorylation status was quantified by calculating the difference in log_2_ coverage ratio +/- RppH between the deletion strain and the WT strain (see panel A). A higher number indicates a larger impact by *mutT4*. **(C)** Transcripts with 5’ ends mapping to previously published TSSs were grouped according to their first two nt. As most transcripts initiate with A or G, those initiating with C or U are not shown. **(D-E)** Transcripts with 5’ ends mapping to published TSSs that initiated with G were binned into quartiles according to the impact of *mutT4* on their 5’ end phosphorylation status. Secondary structure characteristics of the quartile least affected by *mutT4* (Q1) and the quartile most affected by *mutT4* (Q4) were compared. The first 20 nt of each transcript were computationally folded and the probability that the first 5 nt were unpaired in these structures **(D)** as well as the minimum free energy (MFE) of the structures **(E)** were determined. **(F)** The impact of *mutT4* on leadered vs leaderless transcripts was compared. * p ≤ 0.05, ** p ≤ 0.01, **** p ≤ 0.0001. A one-way ANOVA was performed with Tukey’s test for panel C. Mann-Whitney test was performed for panels D, E and F.

We further analyzed the 5’ ends corresponding to TSSs to gain insights into the sequence and secondary structure preference of MutT4 for its mRNA targets. First, we quantified the impact of *mutT4* on each 5’ end by comparing the +/- RppH coverage ratios in the WT strain to those in the *ΔmutT4* strain. We then grouped the 5’ ends based on their first two nucleotides. As most mycobacterial mRNAs initiate with guanosine or adenosine, we focused on A- or G-starting transcripts. The impact of *mutT4* was generally greater for G-starting than A-starting transcripts, and was greatest for those transcripts starting with the dinucleotide GA (**Figure 2 C**).

To increase our sensitivity for detecting relationships between mRNA secondary structure and *mutT4* activity, we divided the G-starting transcripts into quartiles based on the extent to which they were impacted by deletion of *mutT4*. We then compared predicted secondary structure features of the RNAs for Q1, which represents the transcripts with the smallest increase in triphosphorylation when *mutT4* is deleted, and Q4, which represents transcripts with largest increase in triphosphorylation when *mutT4* is deleted. We used two metrics of secondary structure: the MFE of folding for the 5’-most 20 nt of each transcript, and the probability that the first 5 nt in the RNA are unpaired when considering the folding of the first 20 nt. Both metrics indicated that MutT4 has a greater impact on mRNA 5’ ends with less secondary structure (**Figure 2 D** and **E**), suggesting that structured 5’-ends are less permissive for MutT4 binding or catalytic activity.

Previous studies showed that roughly 25% of the Mtb transcripts are leaderless, i.e. they lack a 5’ untranslated region (UTR) and thus a Shine-Dalgarno sequence for ribosome binding (40,60). We wondered if MutT4 had a preference for leaderless vs leadered transcripts and found that indeed the impact of *mutT4* had on average a greater impact on leaderless transcripts (**Figure 2 F**). The difference was subtle but statistically significant. This did not appear to be attributable to the starting nucleotides of the transcripts, because a larger proportion of leadered transcripts than leaderless transcripts were G-starting (62.4% vs 48.8%, respectively; data from (40)).

### Degradation of a subset of transcripts is dependent on pyrophosphate removal by MutT4

Conversion of RNA 5’ triphosphates to monophosphates is considered a rate-determining step for transcript degradation in *E. coli*, where RNase E is a major mRNA degradation protein that is stimulated by binding to monophosphorylated 5’ ends (27,28). Whether this is the case in mycobacteria is an open question; Mtb RNase E was shown to preferentially cleave a monophosphorylated substrate compared to a substrate with a 5’ hydroxyl (61), but to our knowledge, a direct comparison of triphosphorylated vs monophosphorylated substrates has not been reported. RNase J also has important roles in mRNA degradation in some bacteria, and in some of these cases is known to be simulated by 5’ monophosphates (26,29–32), but this has not been tested in mycobacteria. To investigate the relationship between pyrophosphate removal by MutT4 and transcript stability, we correlated the reported mRNA half-lives of Mtb transcripts from a previous study (5) with our own determination of the impact of *mutT4* deletion on the 5’-end phosphorylation state of each transcript. We focused on G-starting transcripts as we found previously that these are preferred substrates for MutT4. We found a significant negative correlation between mRNA half-life and the impact of *mutT4* deletion on 5’-end triphosphate levels (**Figure 3 A**). No correlation was found for A-starting transcripts (**Supplementary Figure 4**). This result suggests that pyrophosphate removal by MutT4 might increase the degradation rates of some transcripts.

**Figure 3.**
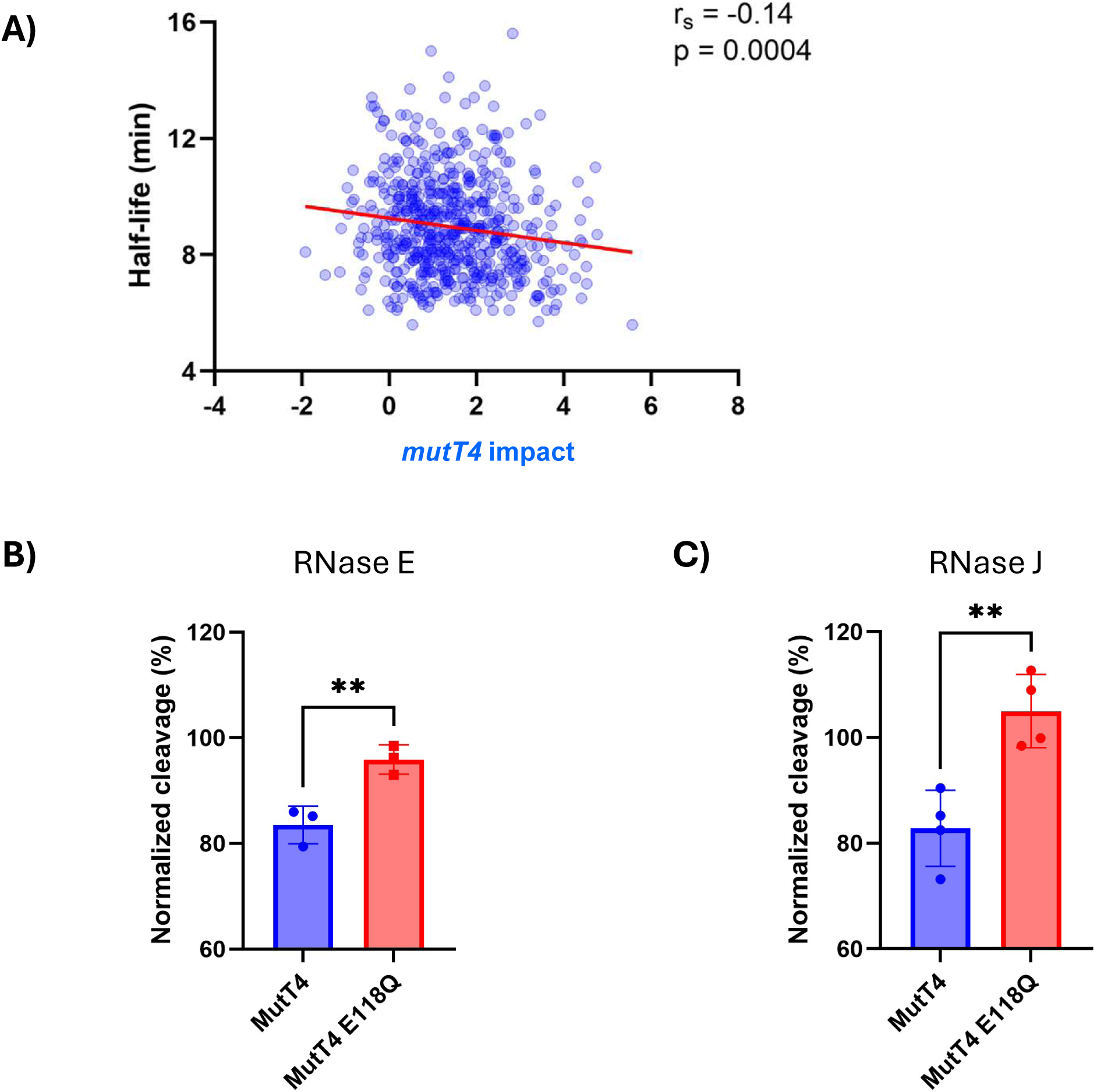
MutT4 activity is associated with faster mRNA degradation *in vivo* and sensitizes transcripts to cleavage by RNase E and RNase J *in vitro*. **(A)** Correlation plot of mRNA half-life (Rustad et al 2013) and impact of *mutT4* on the 5’-end phosphorylation state of G starting transcripts. *mutT4* impact was calculated as shown in Fig 2A. Spearman’s correlation coefficient (r_s_) = -0.14, p-value = 0.0004. 5’ end phosphorylation status was quantified by calculating the difference in log_2_ coverage ratio +/- RppH between the deletion strain and the WT strain. A higher number indicates a larger impact by *mutT4*. (**B-C**) Pre-treatment with MutT4 stimulates degradation of an in vitro transcribed RNA (Rv3248c) by Mtb RNase E (**B**) and RNase J (**C**). Values represent the mean and standard deviation of three or four independent replicates. Cleavage was expressed as percentage decrease in substrate band intensity after incubation with RNase E or RNase J compared to the corresponding untreated control. A representative gel is shown in Supplementary Figure 5. Unpaired t-test was performed for panels B and C. ** p ≤ 0.01

In order to test this hypothesis *in vitro* and investigate the RNases that might be affected by mRNA 5’ end phosphorylation state, we treated *in vitro* transcribed Rv3248c mRNA with purified MutT4, and then tested the ability of Mtb RNase E and RNase J to cleave this mRNA. We found that pretreatment of the mRNA with WT MutT4 (but not the E118Q catalytical-dead version) resulted in increased degradation by both RNase E and RNase J (**Figure 3 B-C** **and Supplementary Figure 5**). This result confirms that mRNA degradation by both RNase E and RNase J is stimulated by 5’ monophosphates in Mtb. Taking this finding together with the major impact of MutT4 on 5’ monophosphate levels in Mtb cells, we conclude that MutT4 is a true constituent of the Mtb mRNA degradation machinery.

### MutT4 forms dynamic biomolecular condensates that appear to localize with RNase E and RNase J

Analysis of the MutT4 protein sequence showed that it has a central folded Nudix domain and intrinsically disordered regions (IDR) at both its N and C-termini (**Supplementary Figure 6**). Interestingly, this protein architecture is similar to that of mycobacterial RNase E, an essential RNase and key component of the mycobacteria mRNA degradation machinery (18). Many IDRs have been reported to be involved in the formation of biomolecular condensates by phase separation through both protein-protein and protein-RNA interactions (reviewed in (62)). Previous studies showed that IDRs are required for the formation of RNase E biomolecular condensates in some bacteria, and we found that *M. smegmatis* RNase E has rifampicin-sensitive punctate localization in cells, a hallmark of phase separation (**Supplementary Figure 7**). The localization of mycobacterial RNase E as dynamic condensates is in agreement with a previous report (63). As the formation of biomolecular condensates is emerging as an important regulatory mechanism in bacteria, we assessed if MutT4 could form this type of structure, using *M. smegmatis* as a model to allow us to do live-cell microscopy. A fusion of the fluorescent protein Dendra2 to the C-terminus of the *M. smegmatis* MutT4 homolog (MutT4_Msmeg_, MSMEG_6927) formed puncta that partially colocalized with RNase E and RNase J condensates (**Figure 4 A**). These results suggest that there could be interactions between MutT4 and RNase E or RNase J. Expression of a MutT4_Mtb_::Dendra2 construct in *M. smegmatis* showed similar localization, indicating that the formation of these structures is conserved across mycobacteria (**Figure 4 B**).

**Figure 4.**
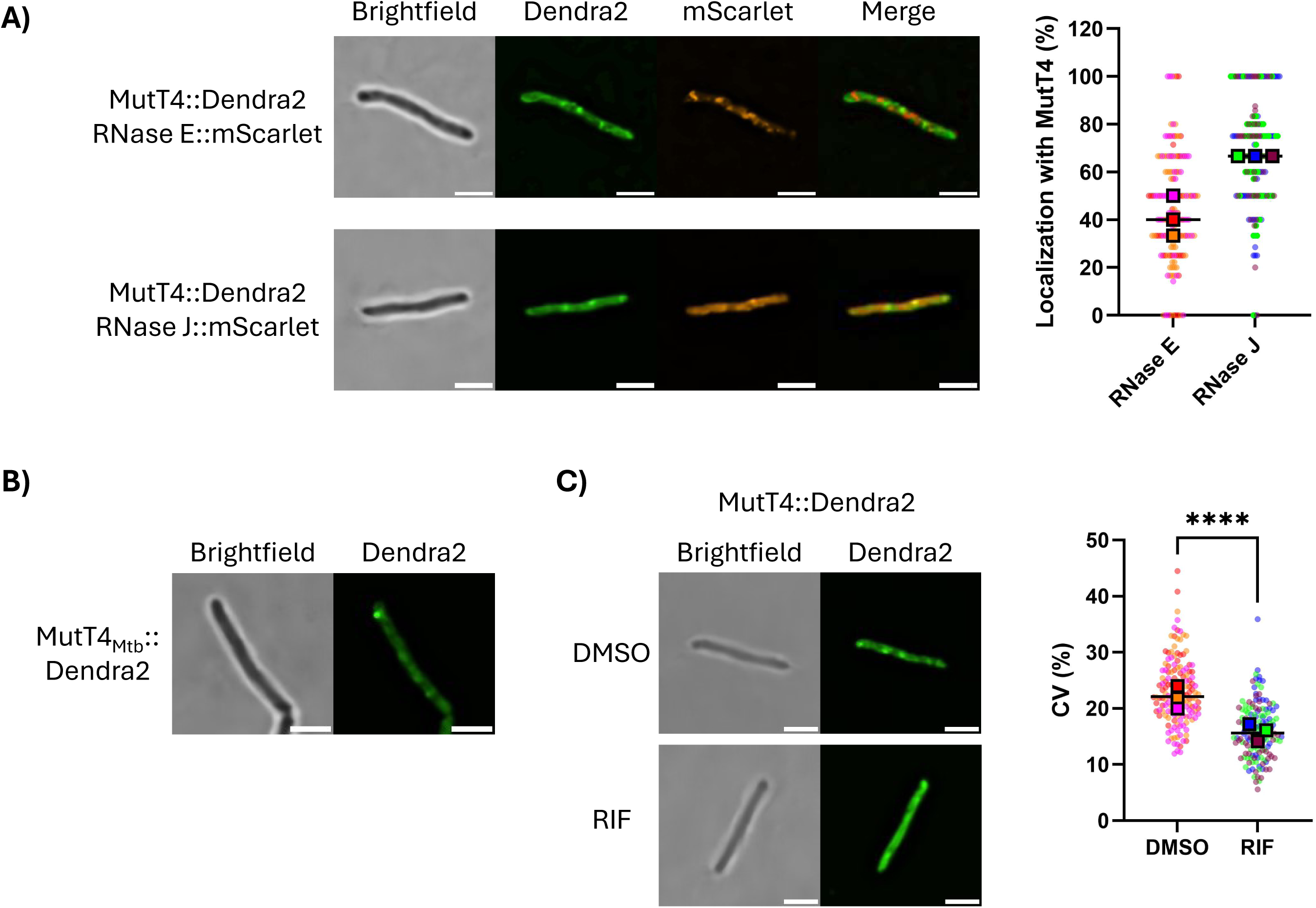
MutT4 forms dynamic biomolecular condensates that appear to localize with RNase E and RNase. **J.** Genes encoding the indicated fluorescent protein fusions were integrated in single copy into phage integration sites in the genome of otherwise WT *M. smegmatis* strain mc^2^155. Live cells were imaged during mid-log phase. All genes are from *M. smegmatis* except in panel B. **(A)** MutT4_Msmeg_ localizes with RNase E and RNase J condensates. Protein localization was analyzed using ImageJ by tracing a line along the center of each bacterium. Fluorescence intensity profiles for the red and green channels were plotted, and overlapping peaks were counted as localizing signals. Percentage localization was calculated as MutT4 localizing peaks divided by total MutT4 peaks per bacterium. **(B)** MutT4_Mtb_ forms condensates when expressed heterologously in *M. smegmatis*. **(C)** Treatment with rifampicin disassembles MutT4_Msmeg_ condensates. *M. smegmatis* expressing MutT4_Msmeg_::Dendra2 was treated with 100 µg/mL rifampicin (RIF) or DMSO as a control for 30 min and imaged. The coefficient of variance (CV) of fluorescence signal within each cell was used as a metric of condensate formation. More punctate signal results in a higher CV. Each dot indicates the CV of fluorescence intensity along a straight line from one cell pole to the other. Scale bar 2 µm for all images. In the plots in A and C, data are from three biological replicates, 50 bacteria each. Medians from each replicate culture are shown as squares and data from individual bacteria as circles. Mann-Whitney test was performed for panel C. **** p ≤ 0.0001

To determine if MutT4 puncta were dynamic condensates affected by the presence of mRNA, we treated *M. smegmatis* with rifampicin to block transcription and thus reduce the bacterial mRNA levels. We found that MutT4 condensates were disassembled in the presence of rifampicin (**Figure 4 C**), confirming that they are dynamic and that they might be RNA-dependent.

### The N-terminal IDR of MutT4 is sufficient for condensate formation

We next examined if the IDRs present in MutT4 participate in condensate formation. To do so, we fused either the MutT4_Msmeg_ N- or C-terminal IDR, or both IDRs, to Dendra2 as shown in the schematic of **Figure 5 A**. As expected, a Dendra2 control showed diffuse cytoplasmatic distribution without the formation of condensates (**Figure 5 B** and **C**). Fusion of MutT4_Msmeg_ N-ter IDR, but not the C-ter IDR, was sufficient for the formation of condensates. Finally, a construct with both IDRs fused to Dendra2 (resembling the domain architecture of MutT4) also formed condensates. This result shows that the N-ter IDR of MutT4_Msmeg_ is sufficient for condensate formation, while the role of the C-ter IDR is unclear.

**Figure 5.**
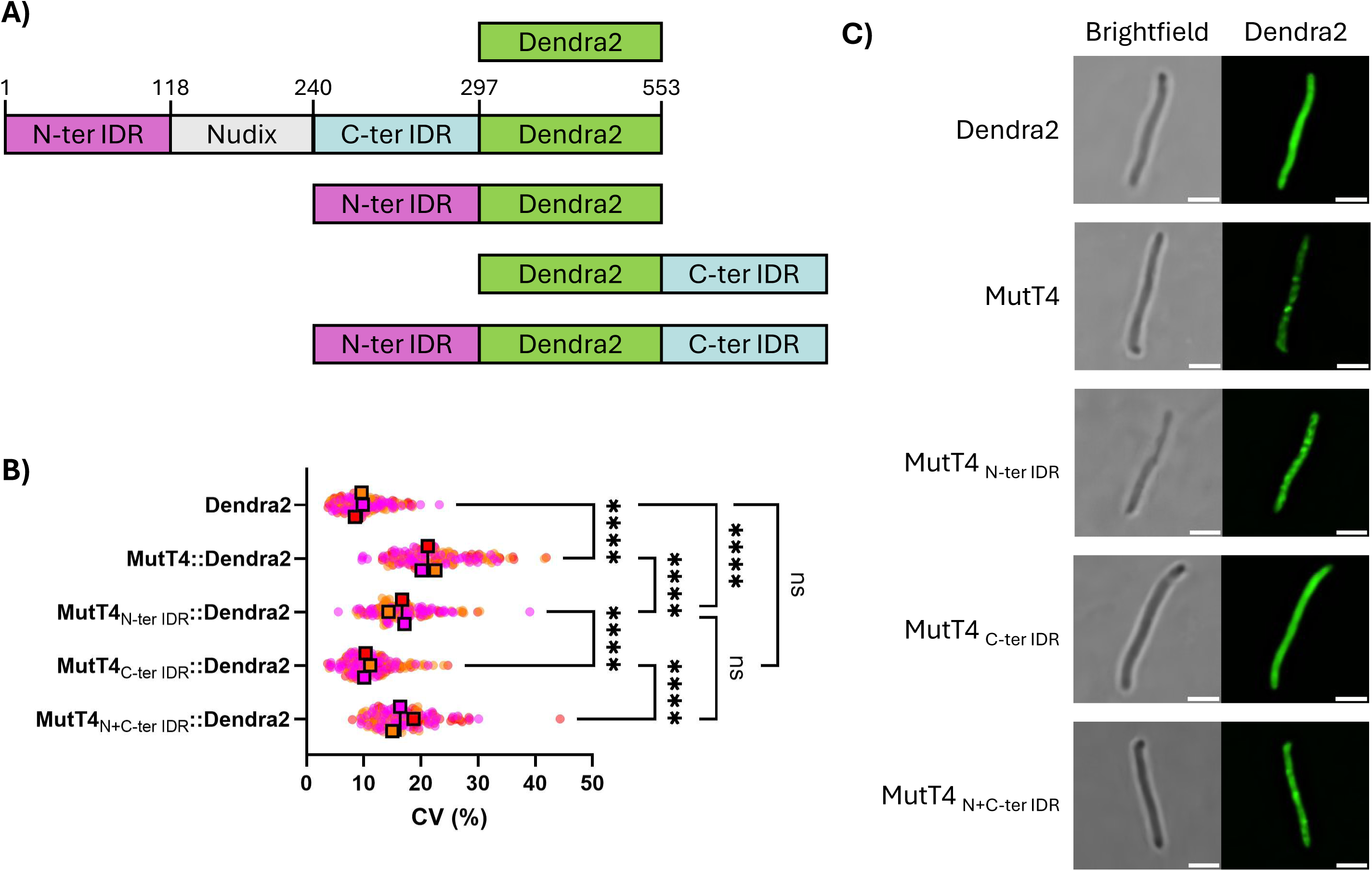
The N-terminal intrinsically disordered region of MutT4_Msmeg_ is sufficient for foci formation. **(A)** Schematic representation of Dendra2 constructs (not to scale). Amino acids are indicated in the MutT4_Msmeg_ full length Dendra2 construct. **(B)** The coefficient of variance (CV) of

To investigate the functional roles of the MutT4 IDRs in Mtb, we complemented Mtb *ΔmutT4* with *mutT4* harboring deletions of the IDRs individually and together. Analysis of the protein expression of these constructs showed that deletion of either IDR decreased the protein abundance, with an additive effect when both IDRs were deleted (**Supplementary Figure 8**). This suggests that the IDRs are involved in the stability of the MutT4 protein in Mtb, and complicates further analysis of their roles since we cannot disentangle the impact of IDR truncation from the impact of altered protein abundance.

### The MutT4 R48G polymorphism found in Lineage 2 has no effect on RNA pyrophosphohydrolase activity or condensate formation

An Arg to Gly point mutation at residue 48 (R48G) in *mutT4* was previously identified in Mtb L2-Beijing isolates (36). As at the time MutT4 was considered to be a DNA repair enzyme based on sequence homology, these studies postulated that the R48G polymorphism could affect the mutation rates of L2 isolates. As our results indicate that Mtb MutT4 is an RNA pyrophosphohydrolase, we sought to analyze if the R48G polymorphism affected this process. We complemented the Mtb *ΔmutT4* mutant strain with either a *mutT4* WT or R48G allele (nucleotide C142G) and used splinted ligation to quantify *in vivo* RNA pyrophosphohydrolase activity on the Rv3248c transcript. We found no significant differences between the strains, indicating that the R48G polymorphism does not affect MutT4 activity (**Supplementary Figure 9 A**). Both of the MutT4 versions were expressed at similar protein levels, indicating that R48G does not change the protein stability (**Supplementary Figure 9 B**). Finally, as the R48G mutation is located in the N-ter IDR, we heterologously expressed MutT4_Mtb_::Dendra2 in *M. smegmatis* and quantified the coefficient of variance (CV) of fluorescence signal within each cell as a metric of condensate formation, and found no significant differences between the WT and R48G alleles (**Supplementary Figure 9 C and D**). Overall, these results indicate that the R48G point mutation does not affect the activity of MutT4.

### MutT4 is not required for Mtb to survive in hypoxia

Adaptation and survival of hypoxic conditions is critical for a successful infection by Mtb (64). We therefore compared the survival of Mtb WT and *ΔmutT4* strains in hypoxic conditions using a modified Wayne model. We detected no differences in the time of methylene blue discoloration (an indirect measurement of oxygen consumption) or the overall viability in hypoxia, indicating that *mutT4* is not involved in adaptation to, or survival of, hypoxia (**Supplementary Figure 10**).

### Absence of MutT4 results in increased vancomycin sensitivity and outer membrane permeability

A previous CRISPRi (65) study indicated that repression of *mutT4* led to increased vancomycin and rifampicin sensitivity. To test these predictions, and determine if they were due to *mutT4* itself and not polar effects on downstream genes, we tested the resistance of Δ*mutT4* to a panel of antibiotics. We found that deletion of *mutT4* increased the sensitivity of Mtb to vancomycin (**Figure 6 A**) and a weak trend in the same direction was seen for rifampicin (**Supplementary Figure 11**). Interestingly, we found no differences in sensitivity to bedaquiline, chloramphenicol, isoniazid, kanamycin, or ofloxacin, suggesting that this increased sensitivity caused by the absence of *mutT4* is specific to vancomycin and rifampicin. As vancomycin and rifampicin have the highest molecular weights of the antibiotics tested, and the site of action of vancomycin is outside of the cell membrane, we hypothesized that the deletion of *mutT4* may increase the permeability of Mtb’s mycomembrane. To evaluate this, we measured the uptake of a fluorescent vancomycin derivative and found that it was increased in the *mutT4* deletion strain (**Figure 6 B**). We then measured mycomembrane permeability in an orthogonal fashion by assessing uptake of an azide-rifamycin conjugate into the Mtb periplasm (**Figure 6 C**) (52,55,66). Uptake of the azide-rifamycin conjugate was greater in the *mutT4* deletion strain than in the WT or complemented strains. Together, these data suggest that the *mutT4* deletion strain has increased mycomembrane permeability.

**Figure 6.**
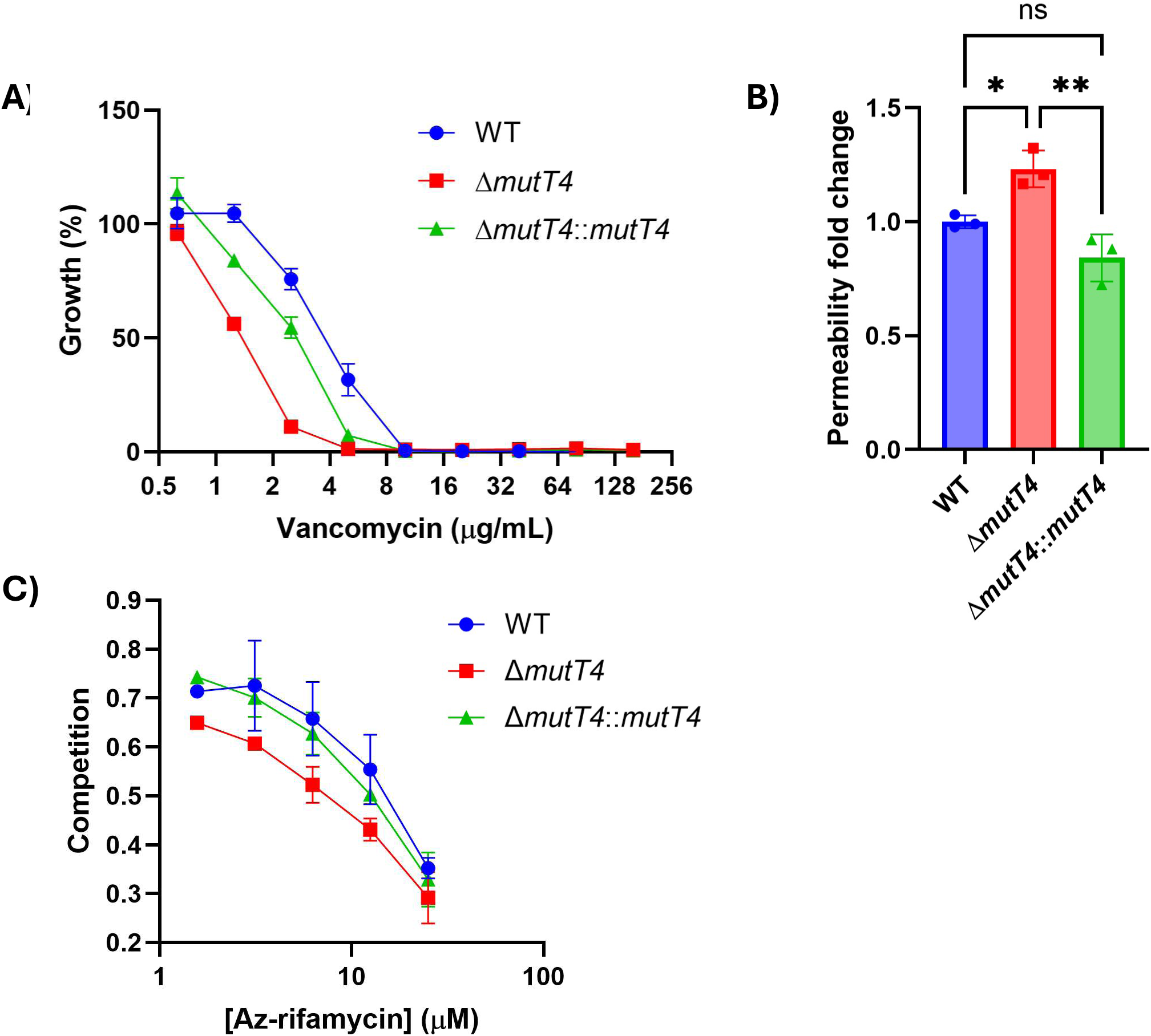
Deletion of *mutT4* leads to increased vancomycin sensitivity and mycomembrane permeability. **(A)** Antibiotic susceptibility testing of Mtb strains against vancomycin. A representative result performed with technical duplicates of three biological replicates is shown. **(B)** Permeability to a fluorescent derivative of vancomycin (FL-VAN). Mtb was incubated with FL-VAN, and permeability to this molecule was measured by flow cytometry and normalized to the WT strain. Higher values indicate higher permeability. Mean and standard deviation of three biological replicates. **(C)**

### Deletion of *mutT4* increases Mtb oxidative stress resistance

In *M. smegmatis*, *mutT4* is required for resistance to oxidative stress (67). To test if this was also true for Mtb, we analyzed resistance to H_2_O_2_ by a disk diffusion assay. Our results show that in contrast to what was previously reported for *M. smegmatis*, deletion of *mutT4* in Mtb increases the resistance to oxidative stress compared to the WT strain (**Figure 7 A**). This difference was reverted by complementation with a WT *mutT4* allele, but not with the E118Q catalytically-dead version, suggesting that RNA pyrophosphohydrolase activity is required for this phenotype. Complementation with the *mutT4* R48G allele showed no difference compared to the *mutT4* WT version. We further confirmed the impact of *mutT4* on H_2_O_2_ sensitivity with an orthogonal approach, by measuring the survival of Mtb after an acute exposure to H_2_O_2_ in 7H9 liquid media. A higher number of viable bacteria were recovered in the *mutT4* deletion strain compared to the WT and complemented strains (**Figure 7 B**).

**Figure 7.**
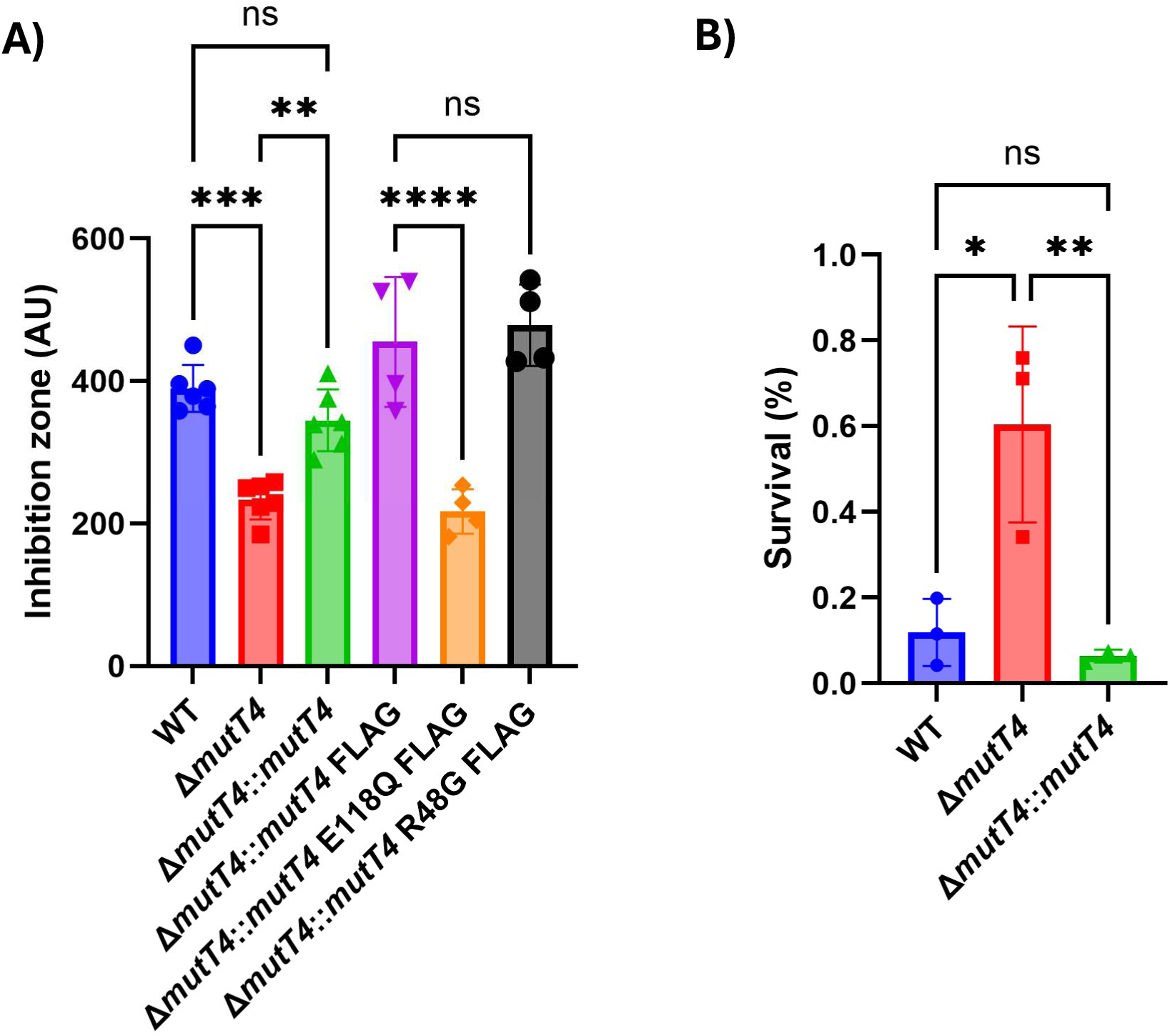
Deletion of *mutT*4 leads to increased resistance to H_2_O_2_ in Mtb. **(A)** Resistance to H_2_O_2_ measured by disk diffusion assay. **(B)** Acute treatment of Mtb with H_2_O_2_ in liquid media. Mtb was exposed to 10 mM H_2_O_2_ for 24 hs and plated to enumerate CFU/mL. Data from three biological replicates are expressed as survival normalized to CFU/mL at time 0. A one-way ANOVA was performed with Tukey’s test for panels A and B. * p ≤ 0.05, ** p ≤ 0.01, *** p ≤ 0.001, **** p ≤ 0.0001

## DISCUSSION

Adaptation to changes in the environment by altering the transcriptome is critical for bacterial survival to stressful conditions. In *E. coli*, RNA degradation can be initiated by RppH pyrophosphate removal and this in turn modulates transcript abundance (25). Here we identify that *mutT4* (Rv3908) encodes the mycobacterial RppH enzyme. While we found that MutT4 processes the 5’ ends of a significant proportion of the transcriptome, integration of our data with those from a previous report of Mtb transcript half-lives (5) suggests that transcript sensitivity to MutT4 is not a major determinant of half-life. Moreover, *mutT4* is a non-essential gene and its deletion does not affect the overall growth rate of Mtb. This result implies that in mycobacteria under optimal growth conditions, 5’-end processing is likely a rate-limiting step for only a subset of transcripts, and that degradation of the majority of transcripts may rely on mechanisms independent of 5’-end chemistry, such as endonucleolytic cleavage by direct entry or 3’-to-5’ exonucleolytic degradation. Alternatively, Mtb could code for other RNA pyrophosphohydrolase(s) that act in coordination with MutT4, not identified by our screening method. The presence of additional enzymes involved in the conversion of RNA 5’ ends from triphosphates to monophosphates other than RppH was also suggested for *E. coli* (68) and *B. subtilis* (26).

Oxidation of dGTP generates the mutagenic nucleotide 8-oxo-dGTP. In *E. coli* the MutT Nudix hydrolase degrades 8-oxo-dGTP, preventing its misincorporation into DNA (69). In a search for mycobacterial genes involved in DNA repair, MutT4 was identified by sequence homology to MutT (36). As Mtb clinical strains of the L2-Beijing lineage carry a point mutation in *mutT4* at codon 48 that results in an Arg to Gly substitution, it was speculated that this variant could lead to an increased mutation rate, a selective advantage for infection and drug resistance (36). However, assigning MutT4 a function as a mutagenic nucleotide cleansing enzyme due to the presence of a Nudix domain is misleading, as this motif is involved in coordination of divalent cation(s) in the active site but does not participate in substrate specificity (70,71). Further, deletion of *mutT4* in Mtb does not lead to an increase in spontaneous mutation frequency (72), suggesting that its role is not related to DNA repair. In *M. smegmatis*, deletion of *mutT4* affects bacterial growth and confers sensitivity to H_2_O_2_ stress and the antibiotics rifampicin and ciprofloxacin (67). All these phenotypes are complemented by expressing *mutT4* of either *M. smegmatis* or Mtb, but not by an *M. smegmatis mutT4* E162A catalytic mutant (equivalent to our Mtb *mutT4* E118Q point mutation). These results confirm that MutT4 serves the same function in both species and that its catalytical activity is essential for this role. Our results here show conclusively that MutT4 is involved in RNA degradation due to its RNA pyrophosphohydrolase activity. We hypothesize that the phenotypes observed by the deletion of *mutT4* in Mtb and *M. smegmatis* are downstream effects due to changes in mRNA 5’-end chemistry that modify transcript stability, and therefore gene expression, during the adaptation to stress. Interestingly, the growth disadvantage caused by the deletion of *mutT4* in *M. smegmatis* implies that this could be the only RNA pyrophosphohydrolase present in this species. RppH in *H. pylori* is also involved in the resistance to oxidative stress, indicating a common link between these two pathways (73). Regarding the MutT4 R48G point mutation in L2-Beijing lineage strains, our results show that this variant has no effect in the RNA pyrophosphohydrolase activity, biomolecular condensates formation, or resistance to oxidative stress related to MutT4. It remains to be determined if this R48G variant has an undetected role in the activity of MutT4 that enhances Mtb pathogenicity, or if it is a result of spontaneous mutation that was expanded clonally.

A significant number of Mtb RNA transcripts are leaderless, as they lack 5’ UTRs (74). This absence of a ribosome binding sequence does not impair their ability to be translated (40,75,76). Previous studies have not, however, considered the possible impact of 5’-end phosphorylation state on translation of leaderless transcripts. It could be the case that changes in the 5’-end chemistry of the mRNA could alter not the transcript stability itself, but its ability to be translated. In this context, the deletion mutant of *mutT4* is a useful model to test this kind of hypothesis, as in this background there is increased triphosphorylation of a large array of mRNAs, including leadered and leaderless transcripts.

Our 5’-end RNAseq results show that MutT4 prefers a G as a first nucleotide and unstructured 5’ ends. *B. subtilis* RppH requires at least two unpaired nucleotides at the 5′ end of its RNA substrates and strongly prefers G as a second nucleotide, while for the first nucleotide A is slightly preferred over G (34,77). Studies on *E. coli* RppH showed that it also requires two unpaired nucleotides at the 5’ end, and has a less strict sequence preference, with A being modestly favored as a first nucleotide (78). Differences in the active site of these enzymes that shape their substrate preference could impact the mRNAs they target, adding a layer of control to tune mRNA levels by selective degradation. Biochemical characterization of RppH orthologs has also showed two distinct mechanisms of catalysis on 5’-triphosphorylated mRNA substrates: while *E. coli* RppH predominantly cleaves a pyrophosphate from the mRNA 5’ end (25), *B. subtilis* RppH cleaves one phosphate at a time (26). Further studies are needed to determine if catalysis by Mtb MutT4 resembles the *E. coli* or *B. subtilis* orthologs.

Unlike *E. coli* and *B. subtilis*, mycobacteria encode for both the endoribonuclease RNase E and the 5’ exo-/endoribonuclease RNase J. *E. coli* RNase E has been shown to prefer 5’ monophosphorylated over triphosphorylated RNA substrates for degradation (27). Studies on RNase J in *B. subtilis*, *Thermus thermophilus*, and *Streptomyces coelicolor* have shown that its exonuclease activity is enhanced by 5’ monophosphates and its endonuclease activity predominates with 5’ triphosphorylated RNA substrates (26,29–32). Structural analysis of RNase E and RNase J support these results by showing the presence of a monophosphate binding pocket, favoring catalysis of these substrates (28,31). To our knowledge, this is the first report that demonstrates an enhanced activity of both Mtb RNase E and RNase J on 5’ monophosphorylated over triphosphorylated RNA substrates. Zeller et al (79) characterized Mtb RNase E with an N-terminal truncation, comparing its activity on RNA substrates with either a 5′ monophosphate or a 5′ hydroxyl group, and found a preference for the 5’ monophosphorylated substrates. Here we extend these results by comparing monophosphorylated and triphosphorylated substrates, and by using full-length RNase E. Endonucleolytic activity has been reported for *M. smegmatis* RNase J on a 5′ triphosphorylated RNA while the 5’ exonuclease activity seems to predominate on 5’ monophosphate substrates (80). Our results confirm that exonucleolytic activity is facilitated by 5’ monophosphates in Mtb as well. We did not observe any clear bands appearing in the reactions with RNase J and fully triphosphorylated substrate, although the full-length substrate band diminished in intensity, suggesting that Mtb RNase J also has substantial exonucleolytic activity on 5’ triphosphorylated substrates. In sum, our results indicate that RNA degradation in mycobacteria can be enhanced by conversion of mRNA 5’ ends to monophosphates.

Our results show that the deletion of *mutT4* in Mtb leads to an increase in vancomycin sensitivity. We further confirmed that the increased vancomycin sensitivity was due at least in part to a disruption of the mycomembrane permeability in the absence of *mutT4*. This result is in line with what has been reported for *E. coli*, as the deletion of *rppH* leads to a higher envelope permeability and increased sensitivity to antibiotics (81). This phenotypic convergence is remarkable given the substantial differences in cell envelope composition between mycobacteria and proteobacteria. The emerging link between RNA metabolism and cell envelope homeostasis is intriguing. MutT4 is coded in an operon that includes the gene encoding the putative lipid II flippase MurJ (Rv3910), which is expected to be required for peptidoglycan biosynthesis. Deletion of *mutT4* appears to increase the mycomembrane permeability, and this phenotype is complemented by an ectopically expressed copy of *mutT4*, indicating that it is unrelated to any possible polar effects on *murJ*. The mechanistic link between MutT4 and cell envelope composition is therefore unknown. Interestingly, RNase J is encoded in an operon together with the gene encoding DapA, a protein involved in biosynthesis of an intermediate for mycobacterial peptidoglycan synthesis (82). *E. coli* RppH activity is stimulated by DapF (83), a metabolic enzyme involved in peptidoglycan biosynthesis (84). Functional links between mRNA degradation and cell envelope biosynthesis therefore seem to be conserved across the bacterial domain. It is tempting to speculate that this functional link could facilitate the rapid changes in bacterial gene expression in response to stressors that impact the cell envelope; more work is needed to investigate this hypothesis.

Biomolecular condensates have recently gained recognition as mechanisms for bacterial cells to organize their cytoplasm into membraneless compartments with distinct roles as well as to stimulate chemical reactions by increasing local concentrations of enzymes and substrates (62,85). Several biomolecular condensates have been described in bacteria, such as BR-bodies (composed of the RNase E-based RNA degradosome and its substrates) (22), RNA polymerase clusters (86), and condensates of the transcription termination factor Rho (87), among others (88). In Mtb, RNase E has been described as forming condensates (63) as well as the ABC transporter Rv1747, which phase separates depending on its phosphorylation status (89). MutT4 condensates could represent sites of active 5’ pyrophosphate removal such as RNA degradation hubs, potentially involving the recruitment of specific mRNAs targeted for degradation into this compartment. It is intriguing that we found a partial colocalization with RNase E condensates, indicating that MutT4 might be recruited by this endonuclease to aid in RNA degradation. Non-colocalizing MutT4 condensates might be RNA degradation hubs that are independent of RNase E, perhaps by recruiting RNase J and/or PNPase. Alternatively, these MutT4 condensates could be a way of sequestering away this enzyme from its mRNA targets to prevent their degradation. Either way, formation of MutT4 condensates in mycobacteria could be another layer of controlling its activity. The determination of the MutT4 protein interactome and if these condensates contain RNA could elucidate their role.

Mtb MutT4 is the only RppH ortholog described so far that has large IDRs, and our results support a model where the N-ter IDR is required for condensate formation. *E. coli* RppH is predicted to have a very short IDR at its C-terminus consisting of only 11 amino acids, and the same is true for the first 10 amino acids of *Bdellovibrio bacteriovorus* RppH. It would be interesting to determine if other RppH orthologs are capable of forming biomolecular condensates, either by an IDR-independent pathway or by being recruited by an IDR-containing protein, as this compartmentalization could be a mechanism that bacteria use for regulating mRNA degradation. Intriguingly, the eukaryotic mRNA decapping enzyme Dcp2 (a Nudix hydrolase) has a large IDR at its C-terminus involved in the formation of P-bodies, a biomolecular condensate enriched in mRNA decay proteins (90). The parallelism of condensate formation by RNA 5’ end modification enzymes (MutT4 in Mtb and Dcp2 in eukaryotes) shows that compartmentalization of this RNA degradation step is broadly conserved.

The results presented here support the designation of MutT4 (Rv3908) as the RppH enzyme in mycobacteria. Further studies on the proteins that interact with MutT4, and the dynamics and role of condensate formation will shed light on this RNA degradation pathway in mycobacteria.

## DATA AVAILABILITY

RNAseq data is available in the NCBI Gene Expression Omnibus (GEO) repository (Accession number GSE292713).

## ACKNOWLEDGEMENTS

We thank members of the Shell lab for helpful discussions. We thank members of the WPI undergraduate lab course BB3527 as well as Edward Chocano Coralla for initial testing of the splinted ligation method. We thank the director of the University of Massachusetts Amherst Flow Cytometry facilities, Dr. Amy Burnside, for her help and advice.

## AUTHOR CONTRIBUTIONS

Conceptualization and Writing (original draft): J.H.C., S.S.S.; Data curation and Software: J.X.; Formal analysis and Visualization: J.H.C., J.X.; Funding acquisition: S.S.S., J.C.S., M.S.S.; Investigation: J.H.C., J.X., I.L., A.R.R., M.R.; Methodology: L.A.R., J.H.C., I.L., J.X., M.S.S.; Writing (review & editing): J.H.C., J.X., I.L., A.R.R., L.A.R., M.S.S., J.C.S., S.S.S.

## FUNDING

This work was supported by the NSF grant number 1652756 (to SSS); NIH grant numbers AI143575 (to SSS and JCS), AI156415 (to SSS), and AI179080 (to MSS); by Welch Foundation grant A-0015 (to JCS); and by the University of Massachusetts Amherst Institute for Applied Life Sciences Midigrant and Core Facilities Incentive Funds (to IL).

## CONFLICT OF INTEREST

None declared.

## SUPPLEMENTAL MATERIALS

Table S1: Primers, plasmids, and strains used in this study.

Table S2: In vitro transcripts used as spike-ins for RNASeq normalization.

Table S3: Mtb H37Rv genes encoding proteins with Nudix domains.

Table S4: RNAseq coverage at transcript 5’ ends corresponding to transcription start sites.

Table S5: RNAseq coverage at transcript 5’ ends corresponding to RNA cleavage sites.

Supplemental Figures 1-11.

## Notes

### Competing Interest Statement

The authors have declared no competing interest.

