## Supplemental Figures for "*Mycobacterium tuberculosis* MutT4 is an RNA pyrophosphohydrolase that forms biomolecular condensates and sensitizes mRNAs to degradation"

### Supplementary Figure 1

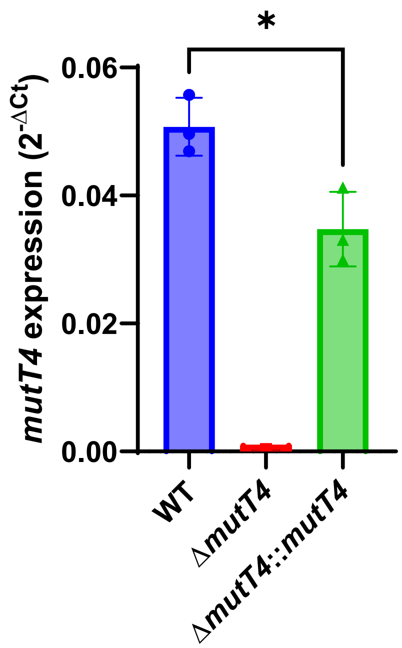


**Supplementary Figure 1.** Comparison of *mutT4* transcript abundance in the WT, deletion, and complemented strains. *mutT4* transcript abundance was normalized to the housekeeping gene *sigA*. Mtb cultures were grown to log phase in 7H9 and the expression of *mutT4* and *sigA* was measured by qPCR. Data from three biological replicates are shown. Unpaired t-test. * $p \leq 0.05$

#### Supplementary Figure 2

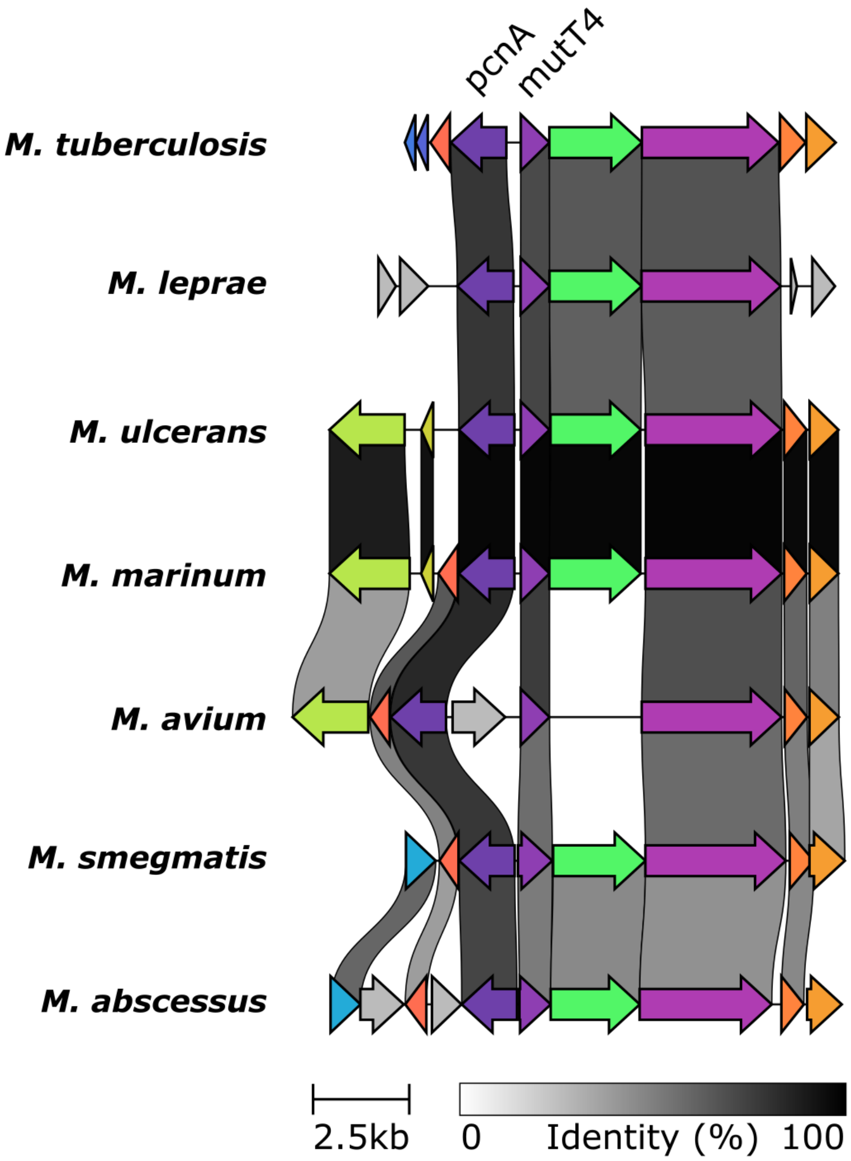


**Supplementary Figure 2.** MutT4 is conserved across mycobacteria in a genomic region that includes the RNA 3'-end CCA-adding enzyme *pcnA*. Figure made with Clinker.

#### Supplementary Figure 3

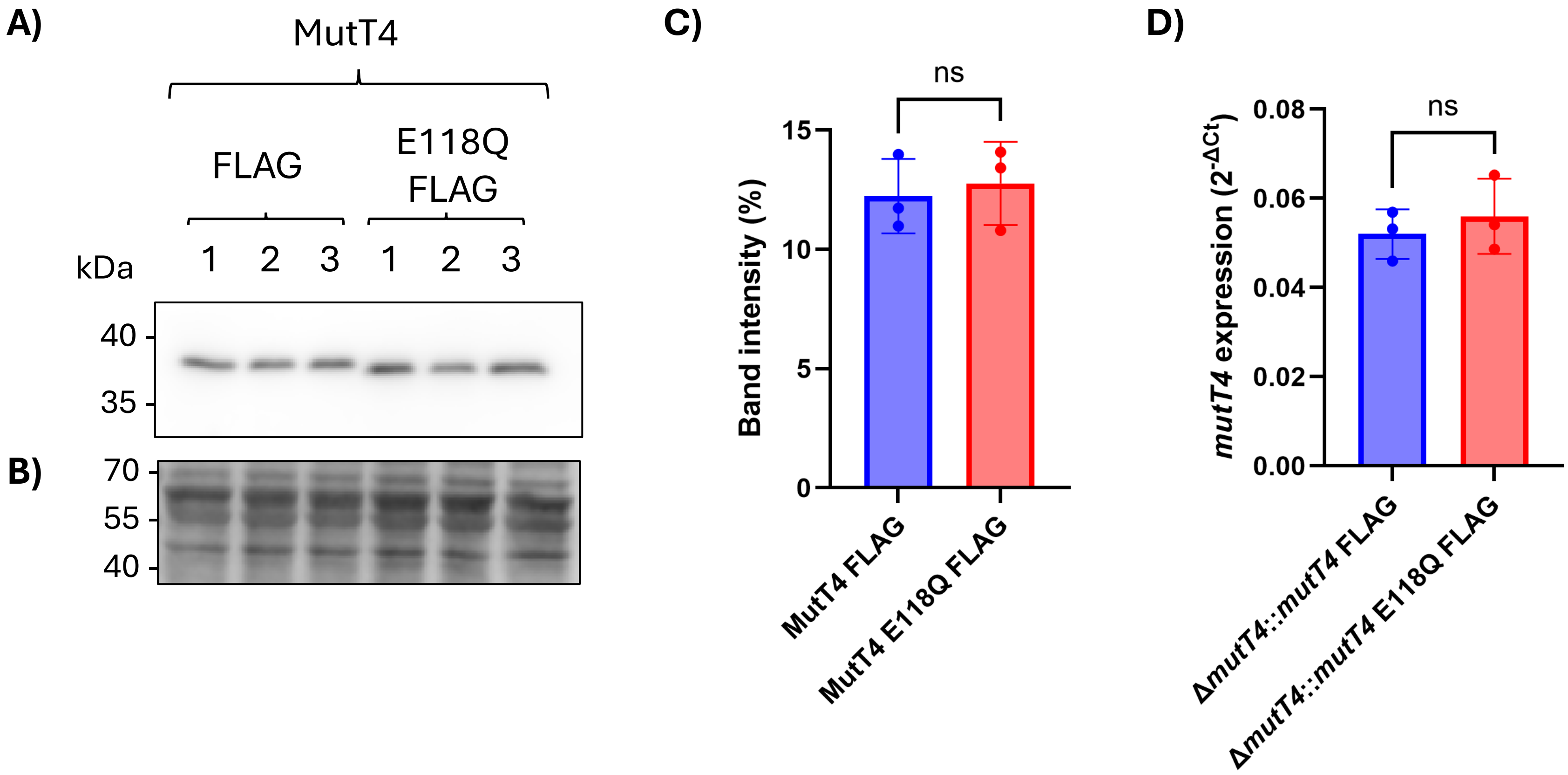


**Supplementary Figure 3.** The E118Q point mutation does not affect MutT4 expression at the protein or RNA level. **(A)** Anti-FLAG Western blot showing the expression levels of FLAG-tagged MutT4 or MutT4 E118Q in the Mtb *ΔmutT4* complemented strains. Each lane represents a biological replicate. **(B)** Proteins on the Western blot membrane were stained with LiCor Revert 700 Total Protein Stain to verify equal loading. **(C)** MutT4 band intensity quantification normalized to the loading control. **(D)** Expression of *mutT4* normalized to the housekeeping gene *sigA*. Mtb cultures were grown to log phase in 7H9 and the expression of *mutT4* and *sigA* was measured by qPCR. Data are from three biological replicates. Unpaired t-test was performed for panels C and D.

**Supplementary Figure 4**

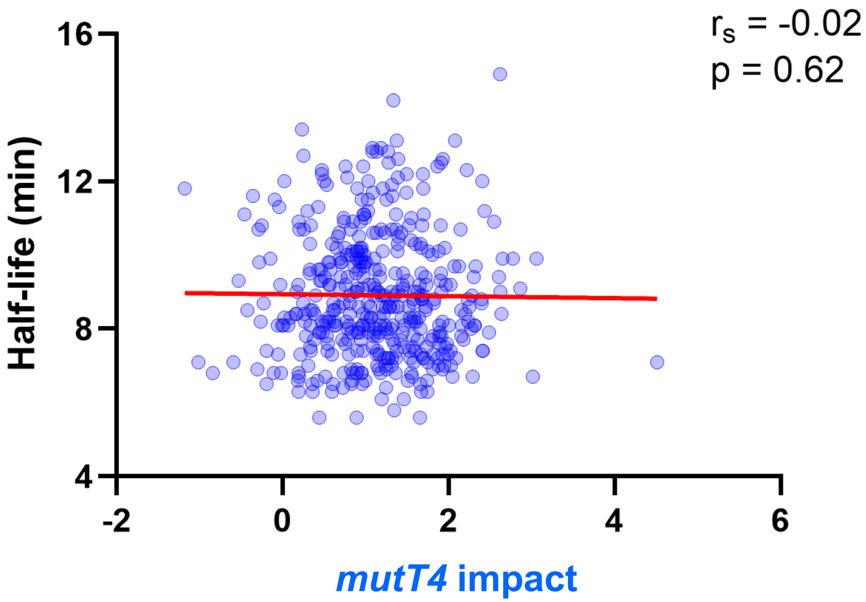


**Supplementary Figure 4.** Correlation plot of mRNA half-life (Rustad et al, 2013) and MutT4 impact on A-starting transcripts. 5' end phosphorylation status was quantified by calculating the difference in $\log_2$ coverage ratio +/- RppH between the deletion strain and the WT strain as shown in Figure 2A. A higher number indicates a larger impact by *mutT4*. Spearman's correlation coefficient ($r_s$) = -0.02, p-value = 0.62.

#### Supplementary Figure 5

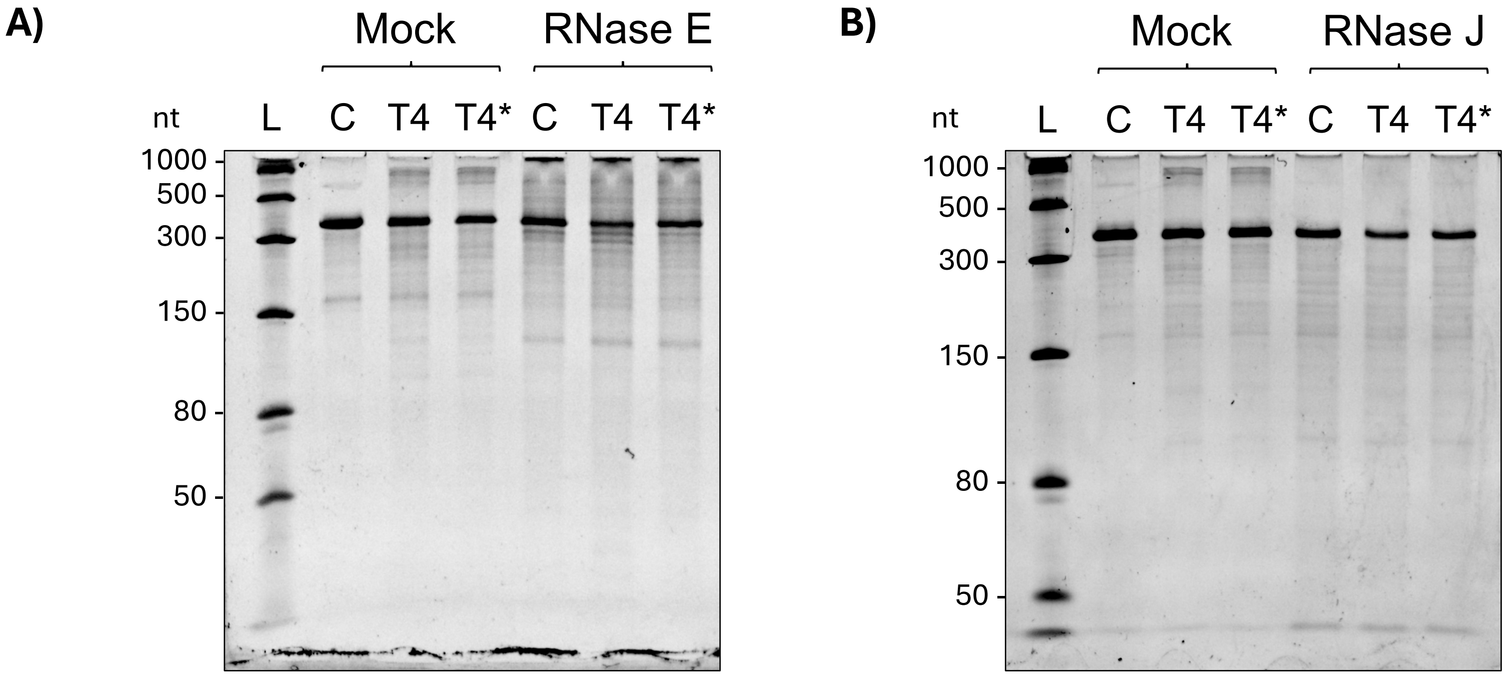


**Supplementary Figure 5.** Representative gels of RNA cleavage assays. In vitro transcribed Rv3248c mRNA was incubated with purified MutT4 (T4), MutT4 E118Q (T4*) or no enzyme (Control, C). Next, mRNA was purified and incubated with no enzyme as a control (Mock), **(A)** Mtb RNase E or **(B)** Mtb RNase J. Samples were separated in a Urea-PAGE gel and stained with SYBR Gold. L, ssRNA ladder.

### Supplementary Figure 6

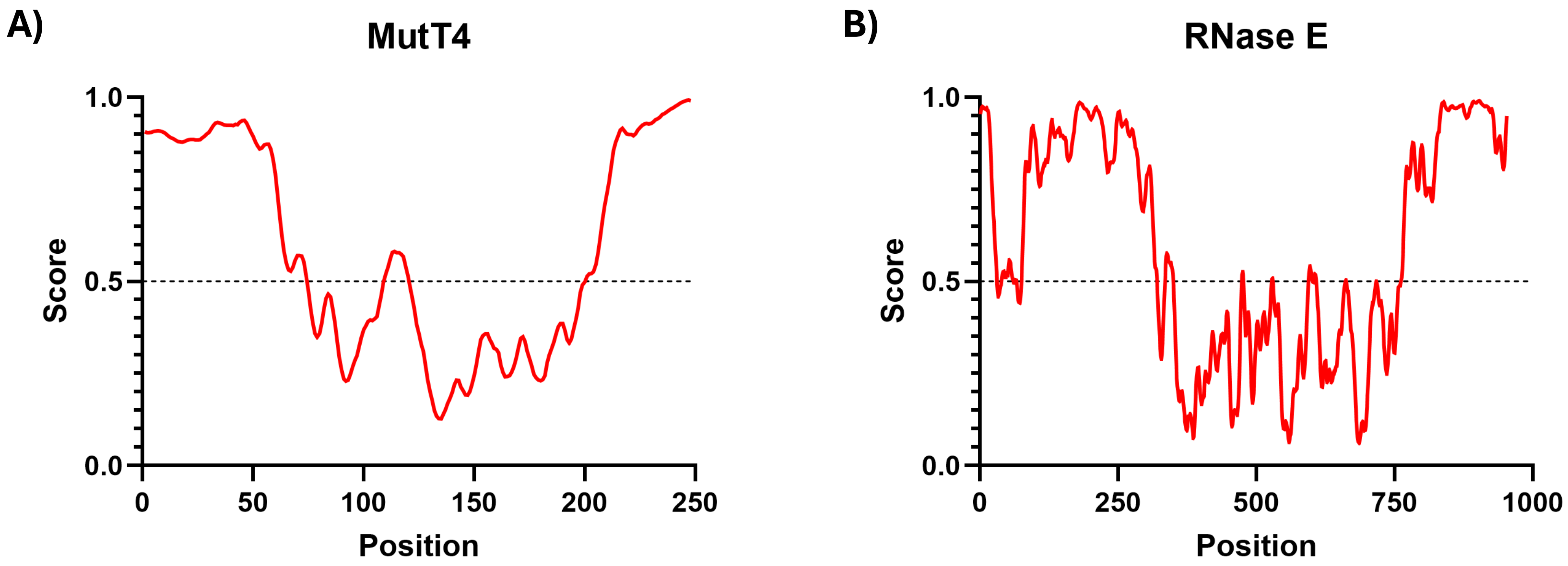


**Supplementary Figure 6.** Intrinsically disordered regions present in the protein sequences of **(A)** Mtb MutT4 and **(B)** Mtb RNase E were identified with IUPred3. Higher values correspond to a higher disorder probability.

#### Supplementary Figure 7

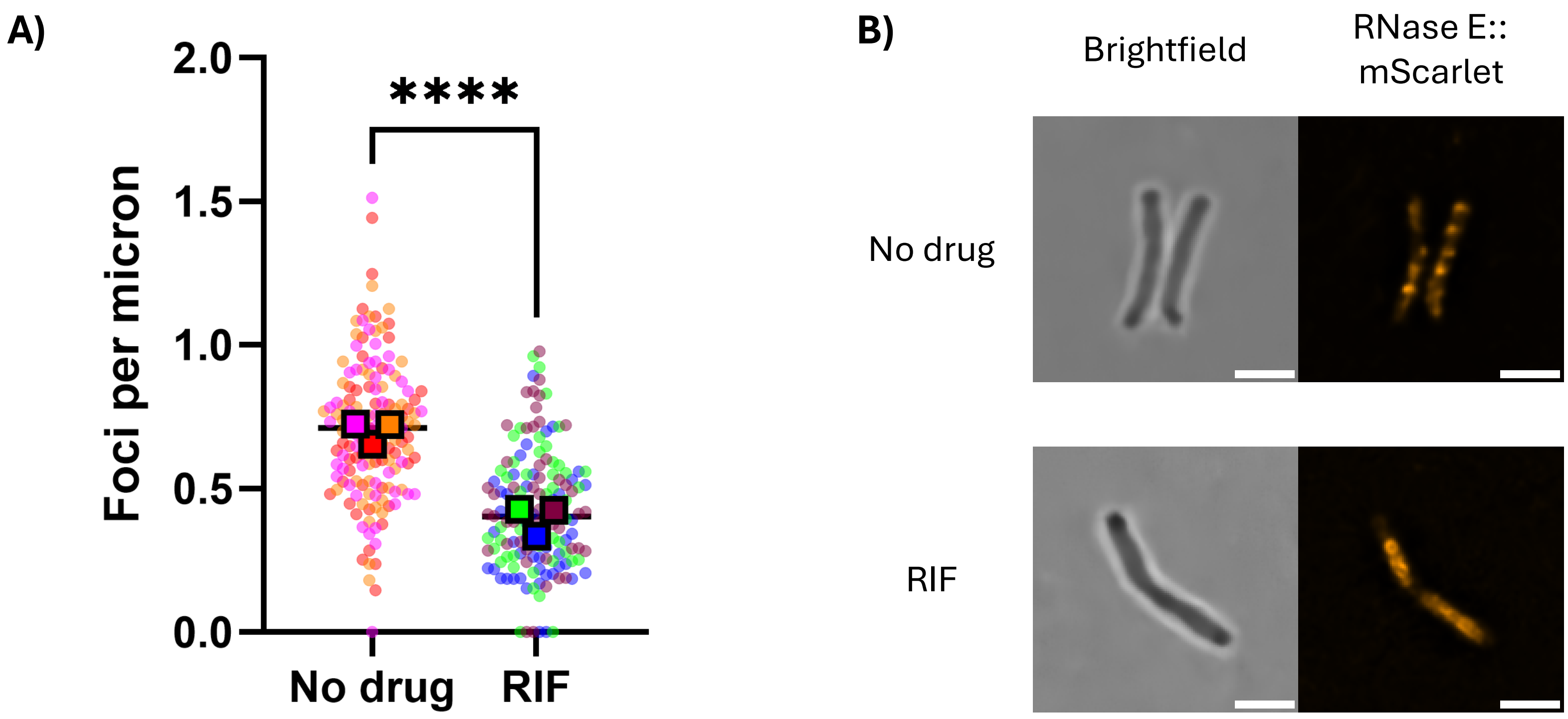


**Supplementary Figure 7.** RNase E forms dynamic condensates in live *M. smegmatis* cells. *M. smegmatis* expressing RNase E::mScarlet was treated with 100 µg/mL rifampicin (RIF) or no drug as a control for 30 min and imaged. **(A)** Treatment with RIF disassembles RNase E condensates. Data from three biological replicates, 50 bacteria each. Medians of the cells in three replicate cultures are shown as squares and data from individual bacteria as circles. **(B)** Representative microscopy images. Scale bar 2 µm. Mann-Whitney test was performed for panel A. **** $p \leq 0.0001$

#### Supplementary Figure 8

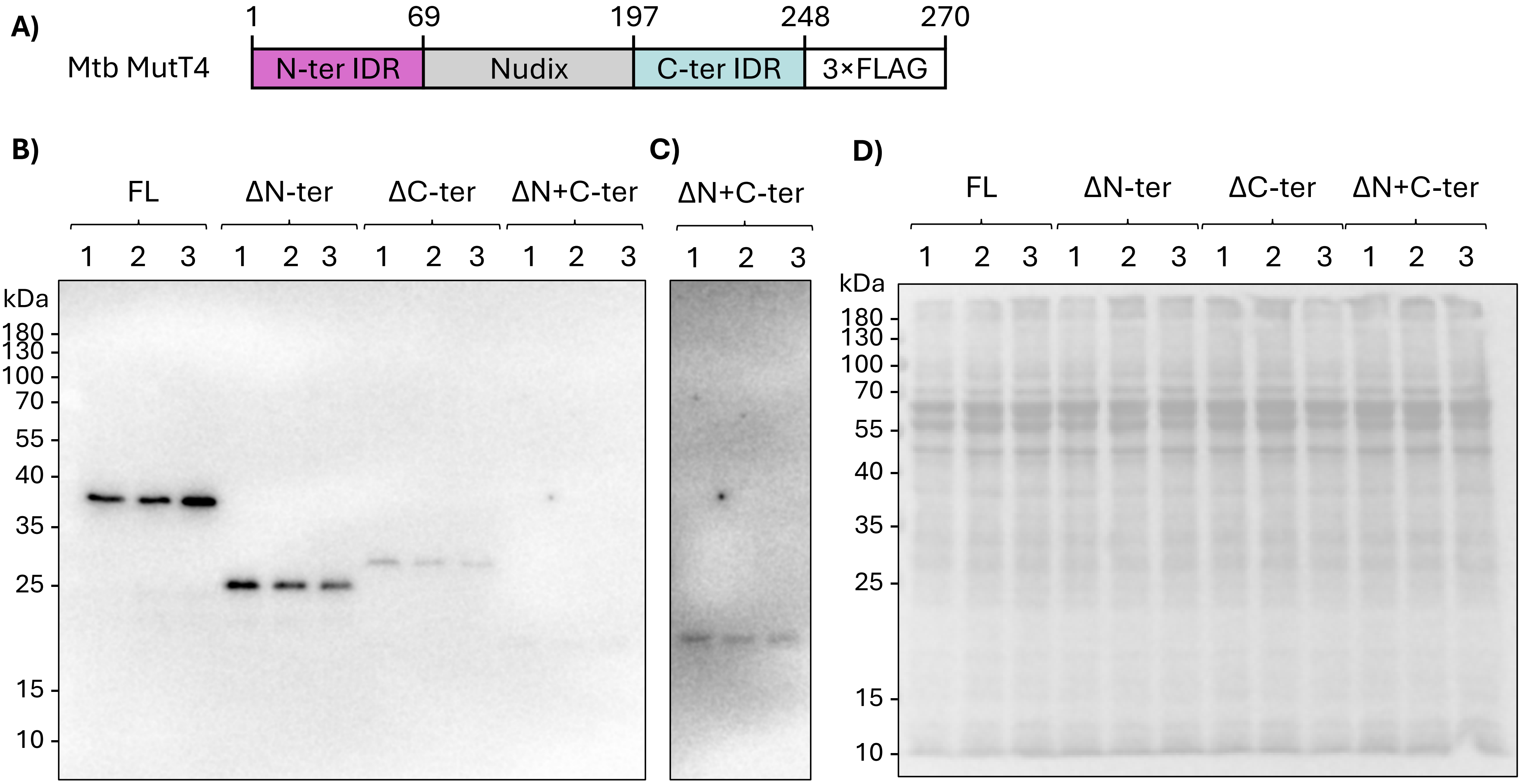


**Supplementary Figure 8.** Deletion of MutT4 IDRs leads to decreased protein levels. **(A)** Schematic of Mtb MutT4 protein. IDR, intrinsically disordered region. Nudix, Nudix domain. Numbers indicate amino acids. Not to scale. **(B)** Mtb protein samples were separated by SDS PAGE and MutT4 detected by anti-FLAG Western blot. Each lane represents a biological replicate. FL, MutT4 FLAG full length. ΔN-ter, MutT4 FLAG with N-ter IDR deletion. ΔC-ter, MutT4 FLAG with C-ter IDR deletion. ΔN+C-ter, MutT4 FLAG with N- and C-ter IDRs deletions. **(C)** Same blot as in B, longer exposure. **(D)** Proteins on the Western blot membrane were stained with LiCor Revert 700 Total Protein Stain to verify equal loading.

#### Supplementary Figure 9

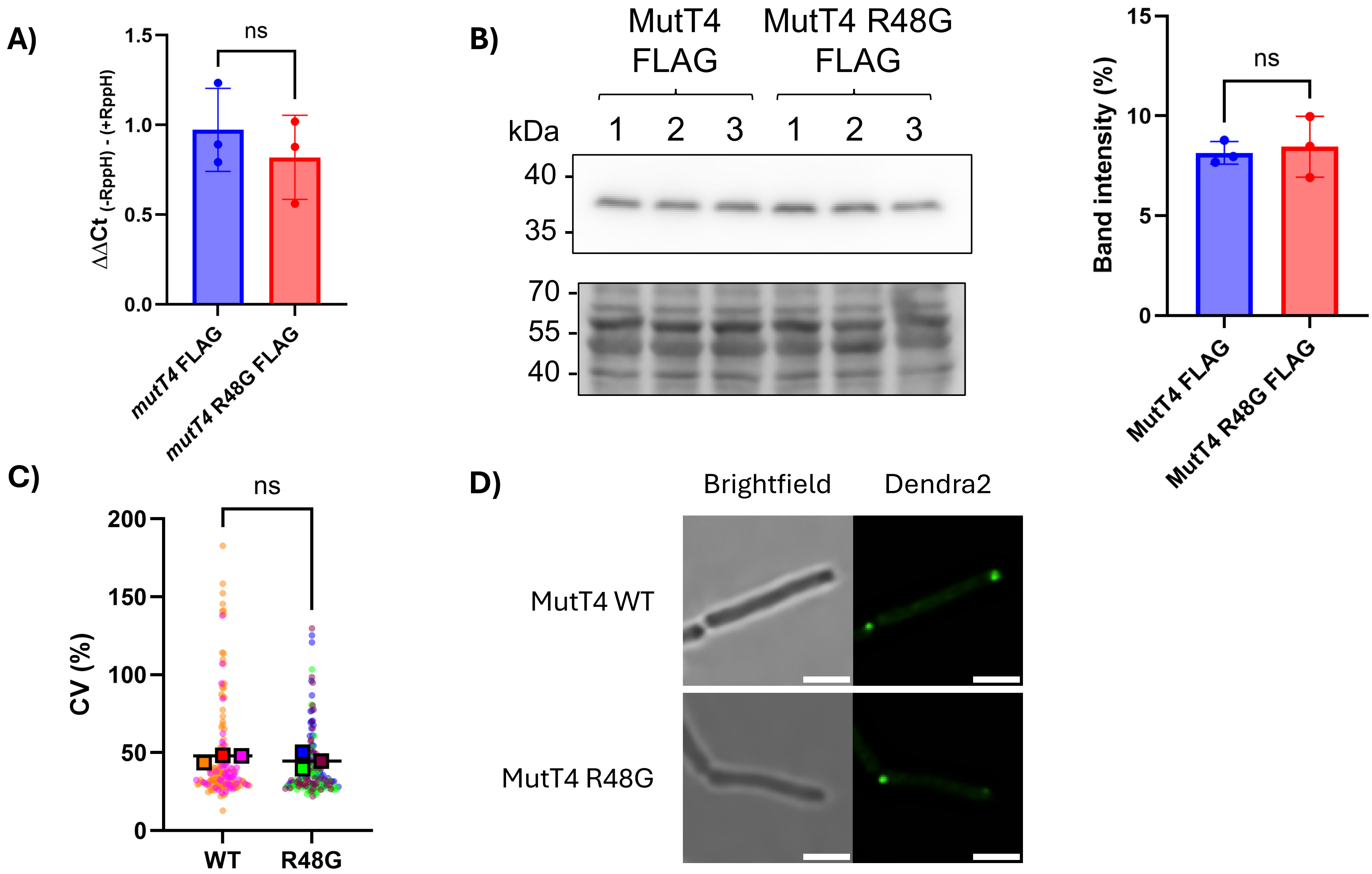


**Supplementary Figure 9.** Functional analysis of the MutT4 R48G variant. **(A)** Measurement of MutT4 *in vivo* activity on target Rv3248c by splinted ligation. Data are from three independent biological replicates. **(B)** Anti-FLAG Western blot (top panel) and band quantification (right panel) normalized to loading control (bottom panel). Samples were from three biological replicates. **(C)** Quantification by confocal microscopy of $MutT4_{Mtb}$::Dendra2 WT or R48G condensates heterologously expressed in *M. smegmatis*. The coefficient of variance (CV) of fluorescence signal within each cell was used as a metric of condensate formation. More punctate signal results in a higher CV. Each dot indicates the CV of fluorescence intensity along a straight line from one cell pole to the other. Scale bar 2 μm for all images. Data from three biological replicates, 50 bacteria each. Medians from each biological replicate are shown as squares. **(D)** Representative confocal microscopy images. Scale bar 2 μm. Unpaired t-test was performed for panels A and B. Mann-Whitney test was performed for panel C.

### Supplementary Figure 10

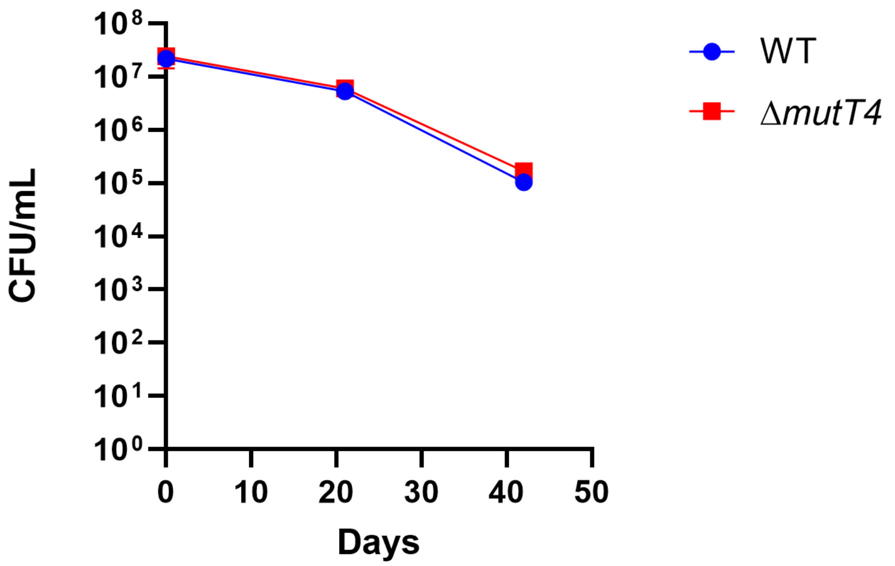


**Supplementary Figure 10.** MutT4 is dispensable for survival in hypoxic conditions. Survival of Mtb WT and *ΔmutT4* strains in hypoxic conditions using a modified Wayne model. Vials were sealed at day 0 to induce hypoxia. Data from three biological replicates.

### Supplementary Figure 11

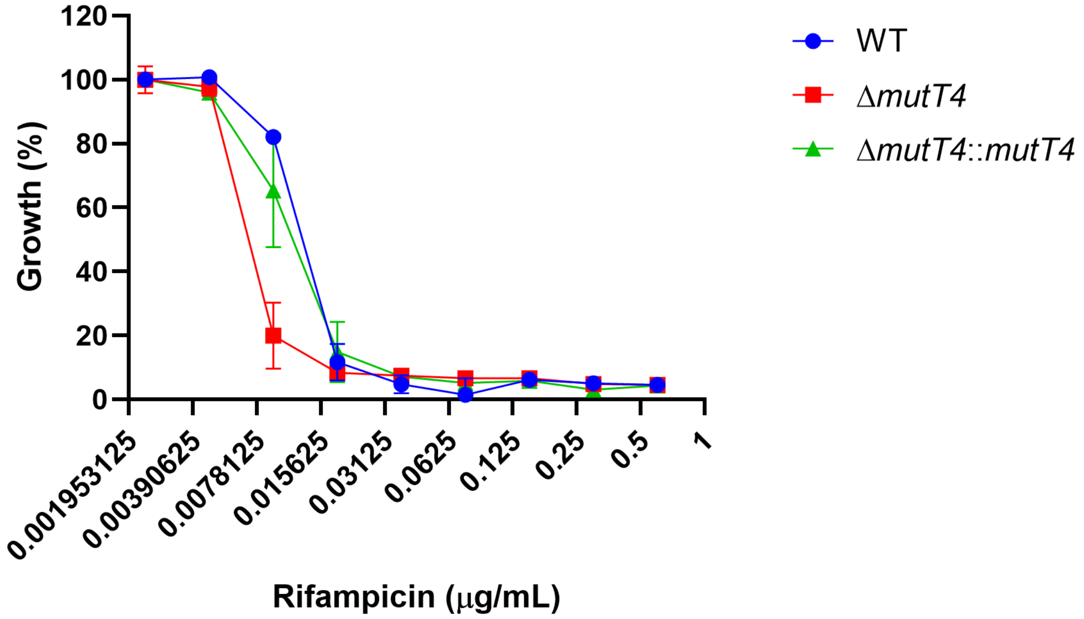


**Supplementary Figure 11.** Deletion of *mutT4* displays a trend towards increased rifampicin sensitivity. Antibiotic susceptibility testing of Mtb strains against rifampicin. A representative result performed in technical duplicates is shown.
